# Genetic analysis of amyotrophic lateral sclerosis identifies contributing pathways and cell types

**DOI:** 10.1101/2020.07.20.211276

**Authors:** Sara Saez-Atienzar, Sara Bandres-Ciga, Rebekah G. Langston, Jonggeol J. Kim, Shing Wan Choi, Regina H. Reynolds, the International ALS Genomics Consortium; ITALSGEN, Yevgeniya Abramzon, Ramita Dewan, Sarah Ahmed, John E. Landers, Ruth Chia, Mina Ryten, Mark R. Cookson, Michael A. Nalls, Adriano Chiò, Bryan J. Traynor

## Abstract

Despite the considerable progress in unraveling the genetic causes of amyotrophic lateral sclerosis (ALS), we do not fully understand the molecular mechanisms underlying the disease. We analyzed genome-wide data involving 78,500 individuals using a polygenic risk score approach to identify the biological pathways and cell types involved in ALS. This data-driven approach identified multiple aspects of the biology underlying the disease that resolved into broader themes, namely *neuron projection morphogenesis, membrane trafficking*, and *signal transduction mediated by ribonucleotides*. We also found that genomic risk in ALS maps consistently to GABAergic cortical interneurons and oligodendrocytes, as confirmed in human single-nucleus RNA-seq data. Using two-sample Mendelian randomization, we nominated five differentially expressed genes (*ATG16L2, ACSL5, MAP1LC3A, PLXNB2*, and *SCFD1*) within the significant pathways as relevant to ALS. We conclude that the disparate genetic etiologies of this fatal neurological disease converge on a smaller number of final common pathways and cell types.

## INTRODUCTION

Amyotrophic lateral sclerosis (ALS, OMIM #105400) is a fatal neurological disease characterized by progressive paralysis that leads to death from respiratory failure typically within three to five years of symptom onset. Approximately 6,000 Americans and 11,000 Europeans die of the condition annually, and the number of ALS cases will increase dramatically over the next two decades, mostly due to aging of the global population (*1*).

Identifying the genes underlying ALS has provided critical insights into the cellular mechanisms leading to neurodegeneration, such as protein homeostasis, cytoskeleton alterations, and RNA metabolism (*2*). Additional efforts based on reductionist and high-throughput cell biology experiments have implicated other pathways, such as endoplasmic reticulum stress (*3*), nucleocytoplasmic transport (*4*), and autophagy defects (*5*). Despite these successes, our knowledge of the biological processes involved in ALS is incomplete, especially for the sporadic form of the disease.

To address this gap in our knowledge, we systematically applied polygenic risk score analysis to a genomic dataset involving 78,500 individuals to distinguish the cellular processes driving ALS. In essence, our polygenic risk score strategy determines whether a particular pathway participates in the pathogenesis of ALS by compiling the effect of multiple genetic variants across all of the genes involved in that pathway. This approach relies solely on genetic information derived from a large cohort and tests all known pathways in a data-driven manner. As such, it provides *prima facie* evidence of the cellular pathways responsible for the disease. Knowledge of the cell types involved in a disease process is an essential step to understanding a disorder. Recognizing this, we extended our computational approach to identify the specific cell types that are involved in ALS. To ensure accessibility, we created an online resource so that the research community can explore the contribution of the various pathways and cell types to ALS risk (https://lng-nia.shinyapps.io/ALS-Pathways/).

## RESULTS

### Pathway analysis used a three-stage study design

Overall, we evaluated the involvement of 7,347 pathways in the pathogenesis of ALS using a polygenic risk score approach (see **Fig. 1A** for the workflow of our analysis). To ensure the accuracy of our results, we divided the available ALS genomic data into three sections. The first of these independent datasets (hereafter known as the reference dataset) was a published genome-wide association study (GWAS) involving 12,577 ALS cases and 23,475 controls (*6*). We used the summary statistics from this reference dataset to define the weights of the risk allele so that greater importance was given to alleles with higher risk estimates.

**Figure 1.**
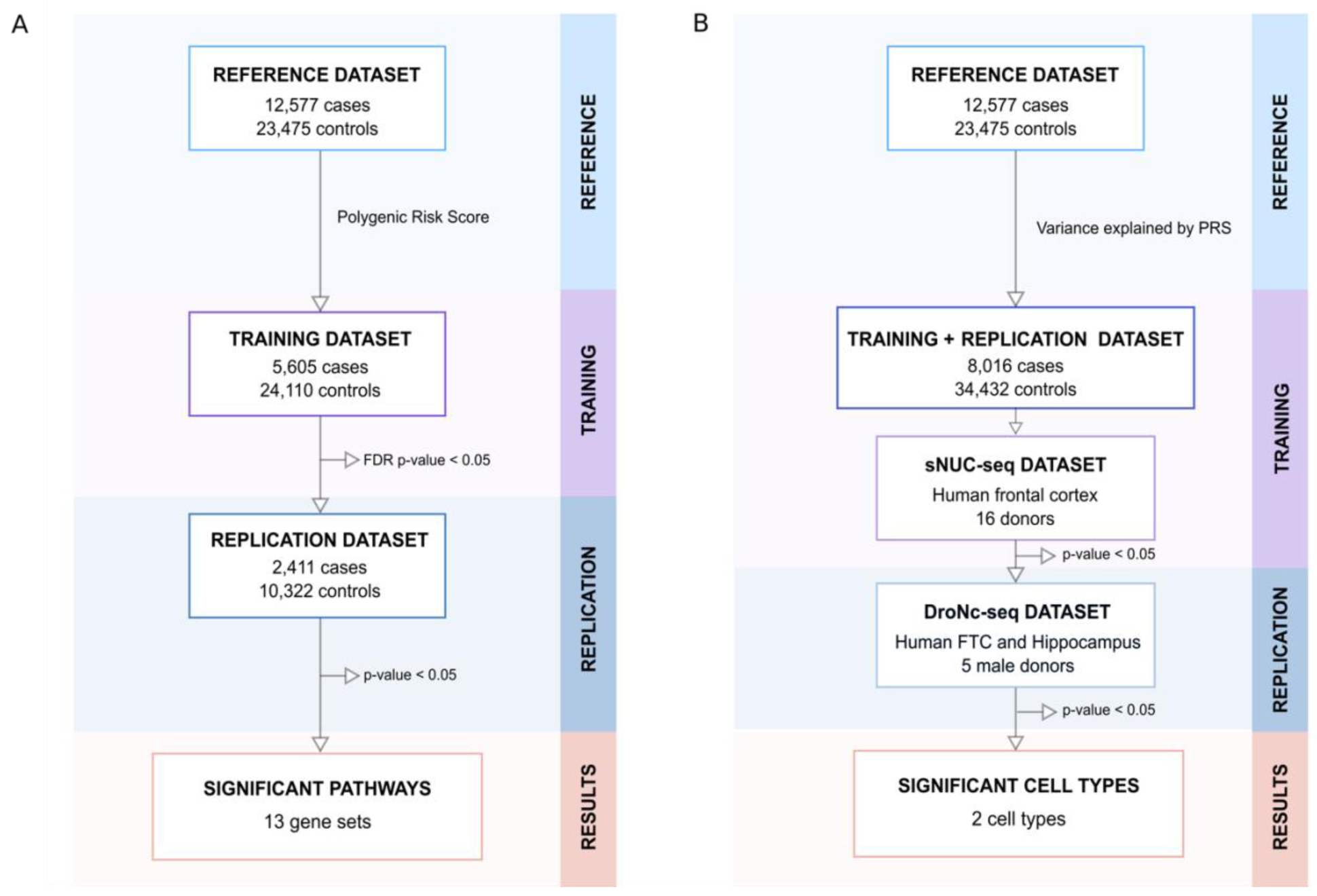
Workflow followed in this study. Polygenic risk score analysis was used to identify (A) biological pathways and (B) cell types contributing to the risk of developing ALS. sNUC dataset was generated in house. DroNc-seq was obtained from Habid et al., 2017. sNUC-seq, single nuclei RNA sequencing; DroNc-seq, droplet single-nucleus RNA-sequencing.

These risk allele weights were then applied to our second dataset (also known as the training dataset) to generate a polygenic risk score estimate for each biological pathway. This training data consisted of individual-level genotype and phenotype data from 5,605 ALS cases and 24,110 control subjects that were genotyped in our laboratory (*7*). We investigated the pathways defined by the Molecular Signatures Database, a compilation of annotated gene sets designed for gene set enrichment and pathway analysis. We focused our efforts on three collections within the Molecular Signatures Database that have been previously validated (*8, 9*). These were the hallmark gene sets (containing 50 pathways), the curated gene sets (1,330 pathways), and the gene ontology gene sets (5,967 pathways).

To ensure the accuracy of our results and control for type I error, we attempted replication of our findings in an independent cohort. For this, we used our third independent dataset (also known as the replication dataset) consisting of individual-level genotype and phenotype data from 2,411 ALS cases and 10,322 controls that were also genotyped in our laboratory (*7*). The pathways that achieved significance in the training dataset (defined as a false discovery rate (FDR)-corrected p-value < 0.05) were selected for replication. Then, we report the pathways that achieved significance in the replication dataset (defined as a raw p-value < 0.05). We estimated the replication cohort’s power to be 98% based on the sample set size, the variance explained by the genetic markers in the training dataset, and the number of independent SNPs with a p-value < 0.05.

We applied a similar polygenic risk score approach to determine which cell types are associated with the ALS disease process (**Fig. 1B**). In essence, a cell type associated with a disease will display a pattern whereby more of the variance of the polygenic risk score is attributable to genes specifically expressed in that cell type. We applied a linear model to detect this pattern in our ALS data, using a p-value of less than 0.05 as the significance threshold. This strategy has become a standard approach for this type of analysis (*10*).

### Biological pathways driving the risk of ALS

We calculated the contribution to ALS risk of 7,347 gene sets and pathways listed in the Molecular Signature Database (***SI Appendix*, Fig. S1**). This genome-wide analysis identified thirteen biological processes, twelve cellular component pathways, and two molecular function pathways with a significant risk associated with ALS in the training data (***SI Appendix*, Table S1**). We independently confirmed a significant association with ALS risk in thirteen of these pathways in our replication cohort. These pathways included (i) seven biological processes, namely *cell morphogenesis involved in neuron differentiation, neuron development, cell morphogenesis involved in differentiation, cell part morphogenesis, cellular component morphogenesis, cell development*, and *cell projection organization* (**Fig. 2A, Table 1)**; (ii) four cellular components, namely *autophagosome, cytoskeleton, nuclear outer membrane endoplasmic reticulum (ER) membrane network*, and *cell projection* (**Fig. 2B, Table 1)**, and (iii) two molecular function terms, namely *ribonucleotide binding* and *protein N-terminus binding* (**Fig. 2C, Table 1**).

**Table 1.**
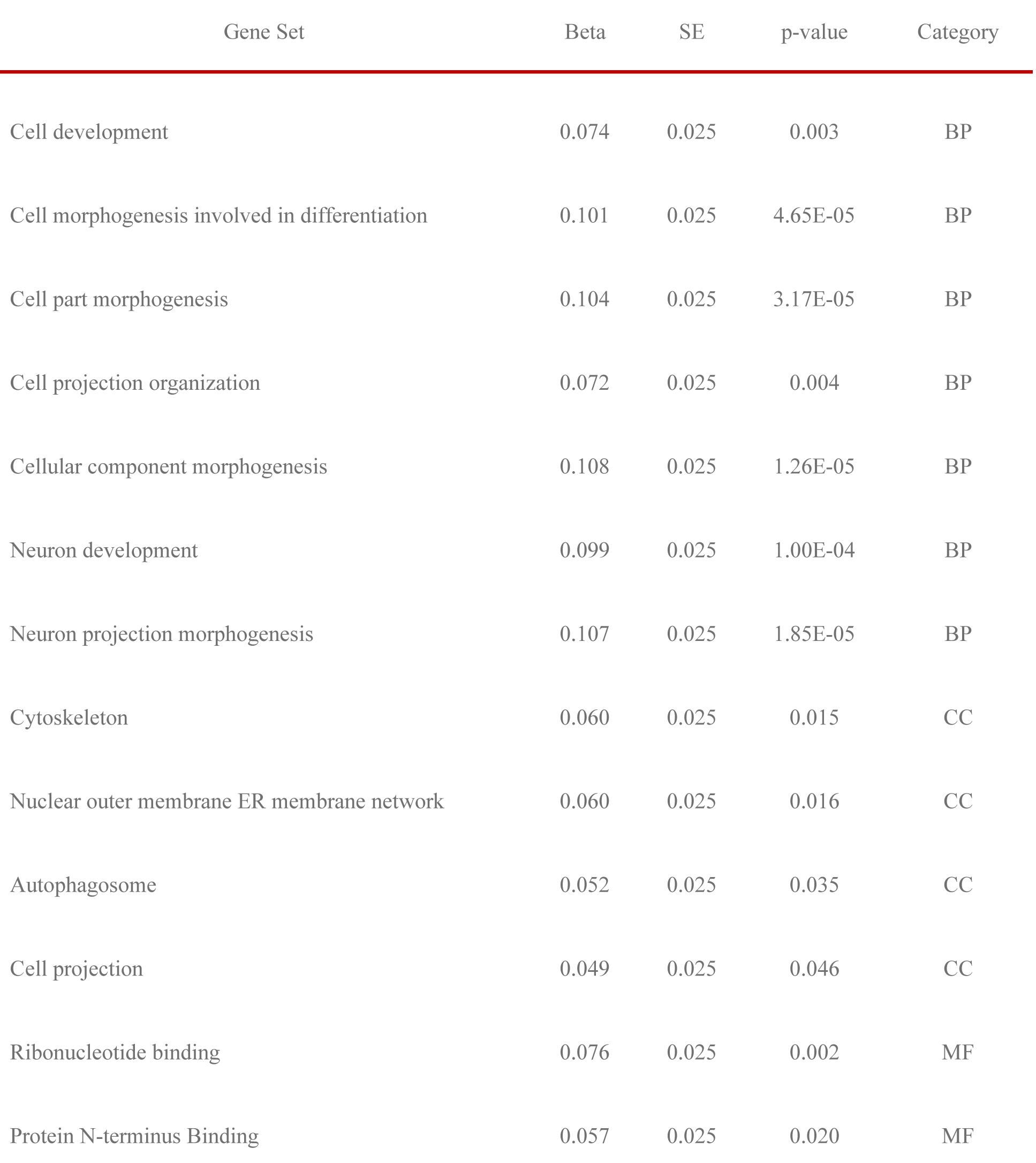
Pathways that were significantly associated with ALS based on polygenic risk score analysis after replication. Beta estimates, standard errors, and p-values are after Z-transformation. SE, standard error; BP, biological process; CC, cellular component; MF, molecular function

| Gene Set | Beta | SE | p-value | Category |
| --- | --- | --- | --- | --- |
| Cell development | 0.074 | 0.025 | 0.003 | BP |
| Cell morphogenesis involved in differentiation | 0.101 | 0.025 | 4.65E-05 | BP |
| Cell part morphogenesis | 0.104 | 0.025 | 3.17E-05 | BP |
| Cell projection organization | 0.072 | 0.025 | 0.004 | BP |
| Cellular component morphogenesis | 0.108 | 0.025 | 1.26E-05 | BP |
| Neuron development | 0.099 | 0.025 | 1.00E-04 | BP |
| Neuron projection morphogenesis | 0.107 | 0.025 | 1.85E-05 | BP |
| Cytoskeleton | 0.060 | 0.025 | 0.015 | CC |
| Nuclear outer membrane ER membrane network | 0.060 | 0.025 | 0.016 | CC |
| Autophagosome | 0.052 | 0.025 | 0.035 | CC |
| Cell projection | 0.049 | 0.025 | 0.046 | CC |
| Ribonucleotide binding | 0.076 | 0.025 | 0.002 | MF |
| Protein N-terminus Binding | 0.057 | 0.025 | 0.020 | MF |

**Figure 2.**
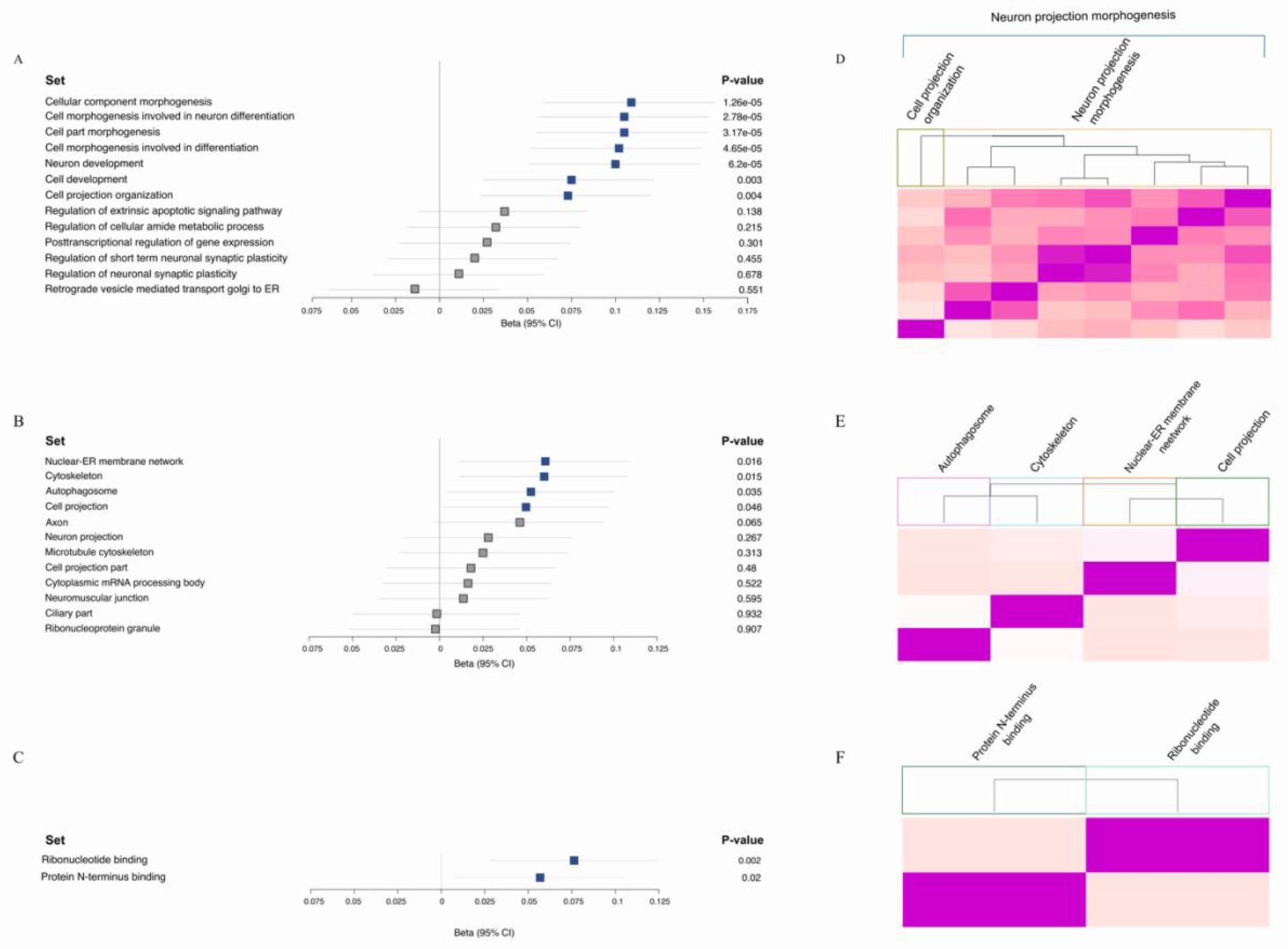
Pathways associated with ALS based on polygenic risk score analysis. The Forest plots show polygenic risk score estimates in the replication cohort for the (A) biological processes, (B) cellular components, and (C) molecular function pathways that were significant in the training cohort. The blue squares represent the significant terms in both the training and replication datasets. The heatmaps depict semantic similarity calculated by *GOSemSim* among the significant (D) biological processes, (E) cellular components, and (F) molecular function. The REVIGO algorithm was used to obtain the cluster representatives. The Forest plot displays the distribution of beta estimates across pathways with the horizontal lines corresponding to 95% confidence intervals. Beta estimates for the polygenetic risk score are after Z transformation.

### Pathways central to ALS risk

There is significant functional overlap among the thirteen pathways that we identified as significantly associated with ALS risk. We sought to more broadly summarize the significant pathways by removing redundant terms. To do this, we computed semantic similarity that is a measure of the relatedness between gene ontology terms based on curated literature. We used the REVIGO algorithm to obtain cluster representatives (*11*). Overall, our results resolved into three central pathways as being involved in the pathogenesis of ALS, namely *neuron projection morphogenesis, membrane trafficking*, and *signal transduction mediated by ribonucleotides* (**Fig. 2D-F, *SI Appendix*, Fig S2)**.

### Pathway analysis among patients carrying the pathogenic *C9orf72* repeat expansion

We found that the *C9orf72* gene was a member of two of our thirteen significant pathways, namely the *autophagosome* and the *cytoskeleton* pathways. We explored whether *C9orf72* was the main driver of these pathways. To do this, we calculated the polygenic risk score associated with these two pathways in *C9orf72* expansion carriers compared to healthy individuals, and non-*C9orf72* carriers compared to healthy individuals. These cohorts consisted of 667 patients diagnosed with ALS who were *C9orf72* expansion carriers, 7,040 patients with ALS who were non-carriers, and 34,232 healthy individuals.

Our analysis revealed that the *cytoskeleton* pathway remained significantly associated with ALS risk in *C9orf72* expansion carriers and non-carriers. This finding indicated that this critical biological process is broadly involved in ALS’s pathogenesis **(Fig. 3B)**. In contrast, only *C9orf72* expansion carriers showed significant risk in the *autophagosome* genes **(Fig. 3A)**, indicating that the *C9orf72* locus mostly drives this pathway’s involvement in the pathogenesis of ALS and points to an autophagy-related mechanism underlying *C9orf72* pathology.

**Figure 3.**
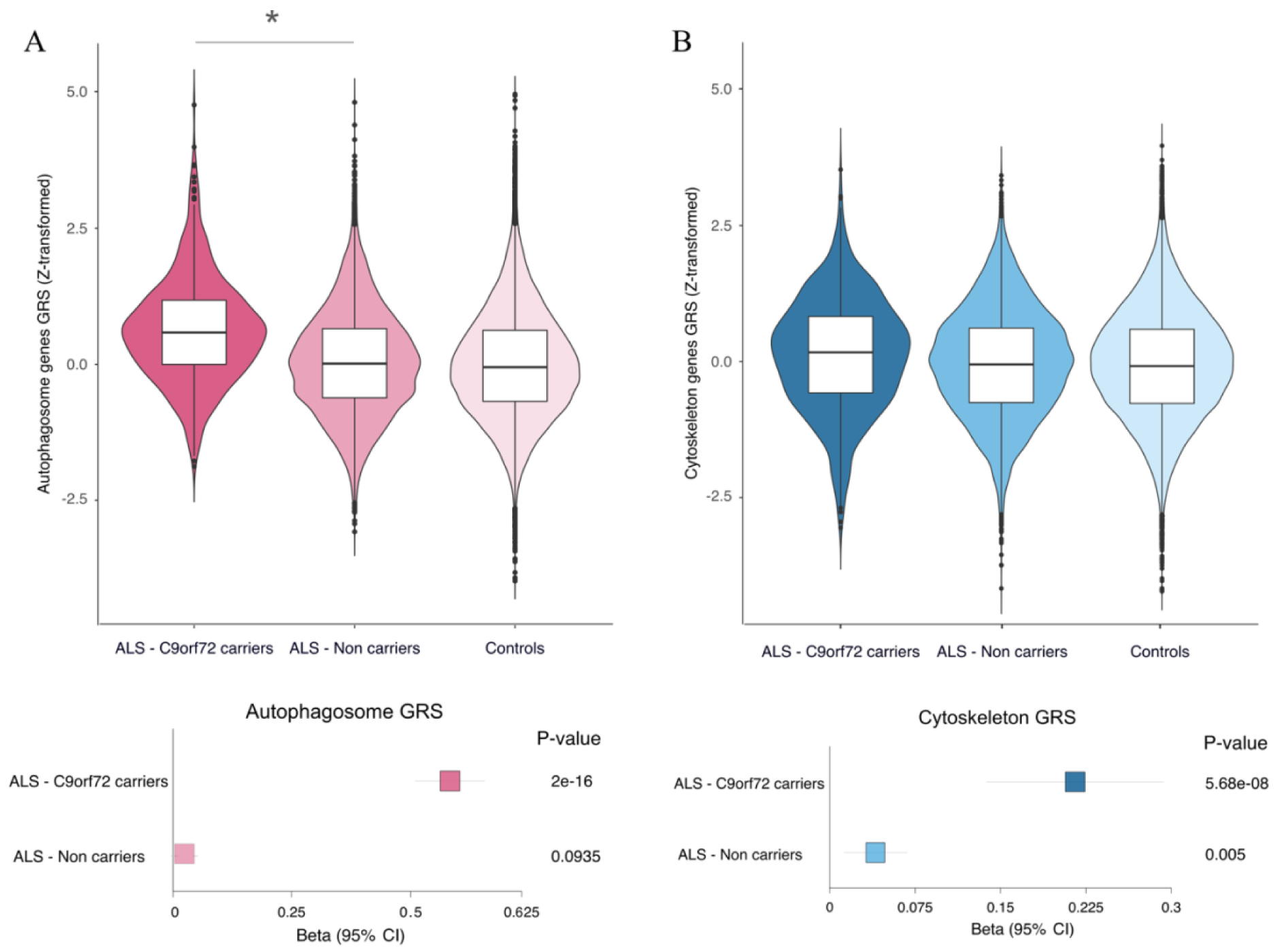
Exploring the role of the *C9orf72* gene in ALS. The polygenic risk scores associated with (A) *autophagosome* and (B) *cytoskeleton* in ALS *C9orf72* expansion carriers (n = 667) compared to healthy subjects (n = 34,232), and ALS non-carriers (n = 7,040) compared to healthy subjects (n = 34,232) are shown in this figure. The upper panels depict the cumulative genetic risk score for each group. The lower panel shows the forest plots of the beta estimates with 95% confidence intervals. Beta estimates are based on the Z-score scale. Genetic risk score mean comparisons from the ALS-non-carriers group compared to the ALS-*C9orf72* carriers via t-test are summarized by asterisks with * denoting a two-sided mean difference at p-value < 0.05.

### ALS polygenic risk is due to genes other than known risk loci

We also examined the contributions of the five genetic risk loci known to be associated with ALS. These loci include *TNIP1, C9orf72, KIF5A, TBK1*, and *UNC13A* (*7*). To do this, we added these five loci as covariates in the analysis of the replication cohort. Our data shows that autophagosome and cell projection were no longer significant. However, the other eleven pathways were still associated, suggesting that there are risk variants contributing to the risk of ALS within these eleven pathways that remain to be discovered (***SI Appendix*, Table S2**).

### Mendelian randomization nominates genes relevant to ALS pathogenesis

Most of the variants associated with a complex trait overlap with expression quantitative trait loci (eQTL), suggesting their involvement in gene expression regulation (*12*). We applied two-sample Mendelian randomization within the thirteen significant pathways (shown in **Table 1**) to integrate summary-level data from a large ALS GWAS (*7*) with data from cis-eQTLs obtained from previous studies in blood (*13*) and brain (*14*)(*15*)(*16*)(*17*). This approach identifies genes whose expression levels are associated with ALS because of a shared causal variant. We used multiple SNPs belonging to the thirteen significant pathways as instruments, gene expression traits as exposure, and the ALS phenotype as the outcome of interest (**Fig. 4A**). Our analyses identified five genes whose altered expression was significantly associated with the risk of developing ALS. These were *ATG16L2, ACSL5, MAP1LC3A, PLXNB2*, and *SCFD1* within blood (***SI Appendix*, Table S3**). Additionally, *SCFD1* was significantly associated with ALS in brain-derived tissue (**Fig. 4B)**

**Figure 4.**
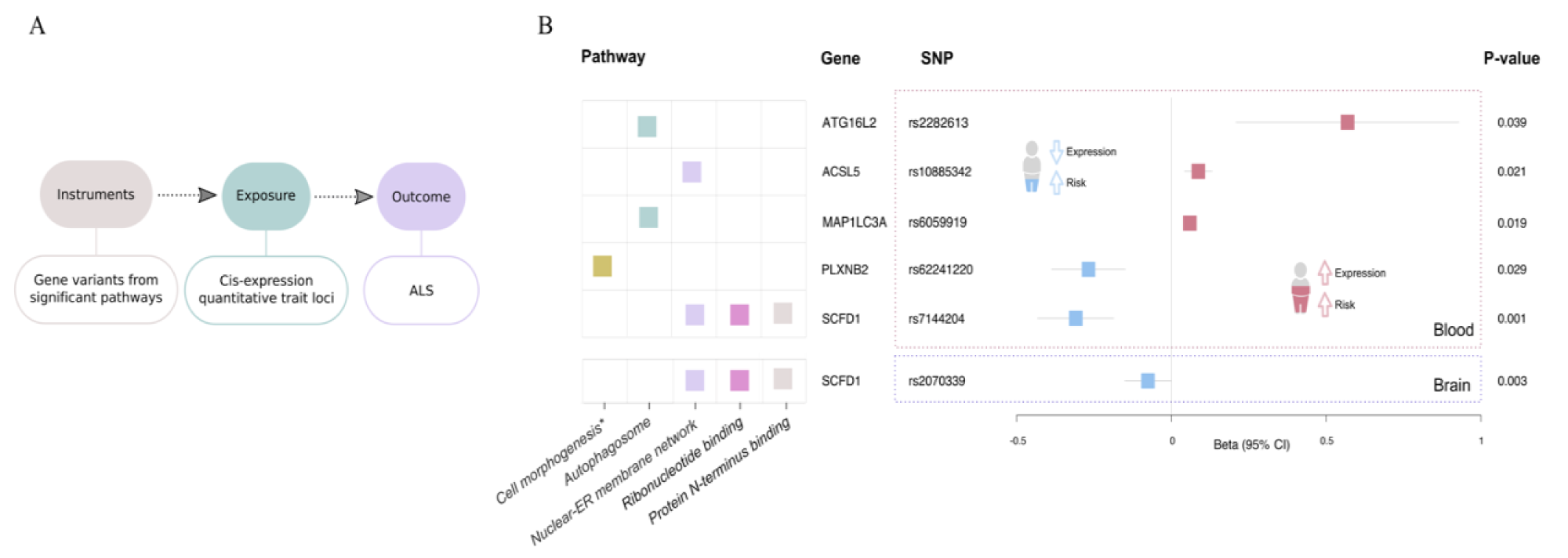
Genes within the significant pathways for which expression was associated with ALS risk based on two-sample Mendelian randomization. (A) A schematic representation of the parameters used for the analysis. (B) The Forest plots display the beta estimates with the 95% confidence intervals shown as horizontal error bars. The grid on the left indicates the pathway to which the gene belongs.

### Cell types involved in the pathogenesis of ALS

We leveraged our large GWAS dataset to determine which cell types participate in the pathological processes of ALS. To do this, we generated a single-nucleus RNA-seq (sNuc-seq) dataset using the human frontal cortex collected from sixteen healthy donors. Each cell was assigned to one of thirty-four specific cell types based on the clustering of the single-nucleus RNA-seq data (**Fig.5 A, B**). We then determined a decile rank of expression for the thirty-four cell types based on the specificity of expression. For instance, the *TREM2* gene is highly expressed only in microglia. Thus, the specificity value of *TREM2* in microglia is close to 1 (0.87), and it is assigned to the tenth decile for this cell type. In contrast, the *POLR1C* gene is expressed widely across tissues. Consequently, it has a specificity value of 0.007, and it is assigned to the fourth decile of microglia and a similar low decile across other cell types.

The premise of this type of analysis is that, for a cell type associated with a disease, more of the variance explained by the polygenic risk score estimates will be attributable to the genes more highly expressed in that cell type. To test this hypothesis, we applied linear regression models to detect a trend of increased variance with the top deciles, a pattern indicating that a particular cell type is involved in the pathogenesis of ALS (*10*). This approach identified two subtypes of *cortical GABAergic interneurons* (*PVALB* and *TOX*-expressing neurons; and *ADARB2* and *RELN*-expressing neurons) and *oligodendrocytes* (*OPALIN, FCHSD2*, and *LAMA2 -*expressing oligodendrocytes) as associated with ALS risk (**Fig. 5, C**).

**Figure 5.**
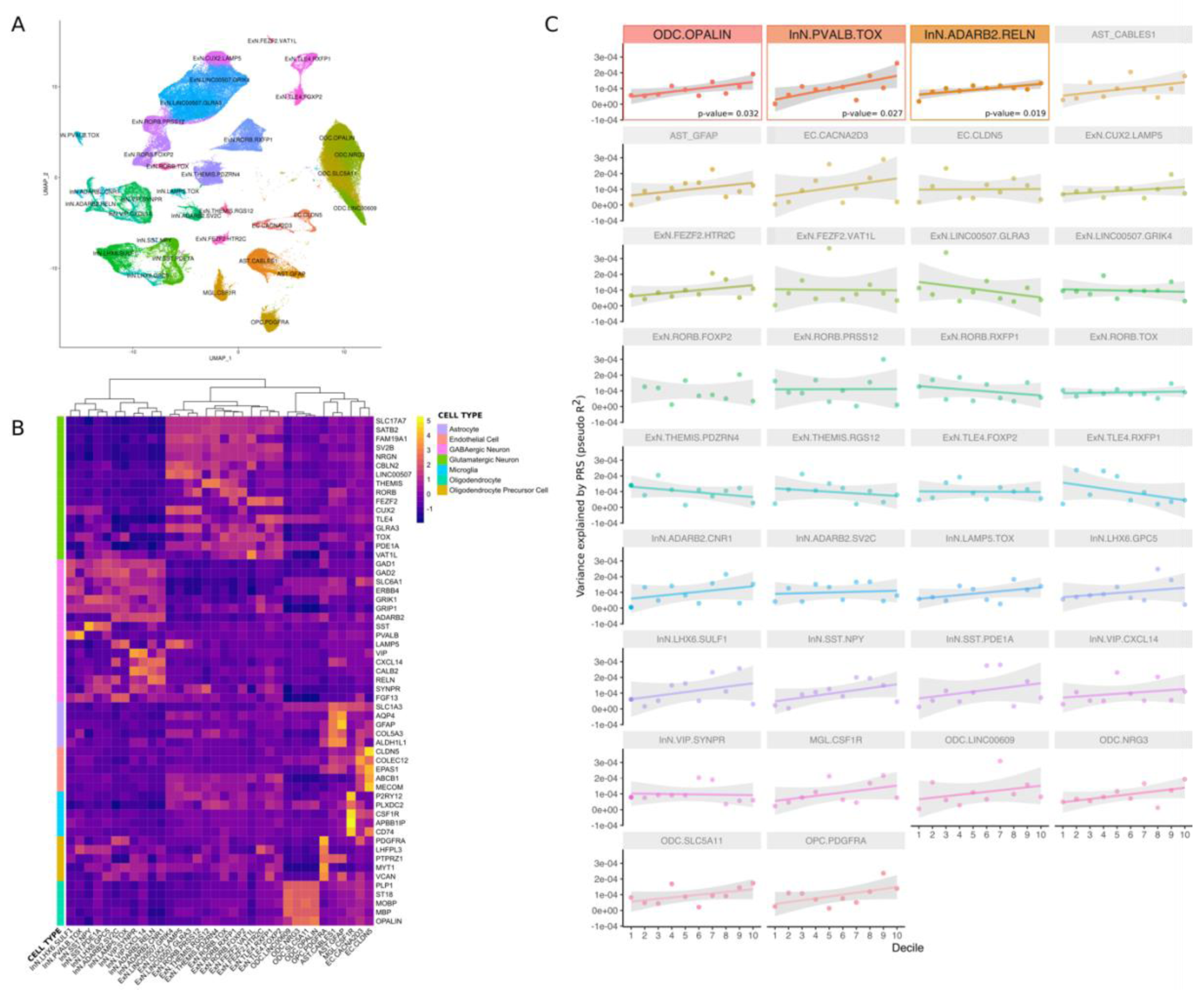
The phenotypic variance explained by polygenic risk score across human cortex cell types. (A) Unsupervised UMAP clustering identifies thirty-four cell types in the human cortex. (B) Heatmap representing the gene expression per cluster. (C) The y-axis corresponds to the phenotypic variance explained by the polygenic risk score (pseudo-R_2_), and the x-axis depicts deciles 1 to 10. The color pictures show the significant cell types and the p-values of the linear regression fit models. The semitransparent pictures show the cell types that were not significantly associated with the disease. The regression line depicts the association between PRS.R2 (pseudo R2, adjusted by prevalence) and the specificity decile in each cell type. The grey shading shows the 95% confidence interval of the regression model. Astrocyte (AST), endothelial cell (EC), excitatory neuron (ExN), inhibitory neuron (InN), microglia (MGL), oligodendrocyte (ODC), oligodendrocyte precursor (OPC).

To confirm these findings, we used an independent dataset consisting of droplet single-nucleus RNA-seq (DroNc-seq) from the human cortex and hippocampus obtained from five healthy donors (*18*). Our modeling in this second human dataset confirmed our previous findings: *PVALB*-expressing *GABAergic neurons* and *oligodendrocytes* were significantly enriched in ALS risk (see ***SI Appendix*, Fig. S3**).

Finally, we attempted to replicate our findings in a well-validated dataset based on single-cell RNA-seq (scRNA-seq) data obtained from mouse brain regions (*10*). The advantage of this nonhuman dataset is that it is based on single-cell RNA-seq, a difficult technique to apply to human neurons, but which captures transcripts missed by single-nucleus RNA-seq that may be important for neurological disease (*10*). Like the human data, cortical parvalbuminergic interneurons again showed enrichment in ALS risk using the mouse dataset (see ***SI Appendix*, Fig. S4**). Oligodendrocytes were not significantly associated with ALS in the mouse superset. However, this is likely related to the depletion of the oligodendroglia lineage in the single-cell RNA-seq data, a known issue with this technology(*10*).

## DISCUSSION

A striking aspect of our analysis is that it identified a relatively small number of biological pathways as central to the pathogenesis of ALS. Considering the clinicopathological and genetic heterogeneity across ALS, the finding of such a small quantity of universal themes is unexpected. Our results illustrate how multiple unrelated genetic causes can lead to a similar downstream outcome, namely motor neuron degeneration. Unraveling how disruption of these three fundamental biological processes predisposes to ALS may yield therapeutic targets that are effective across all patients with ALS.

The importance of *membrane trafficking* in ALS has been widely reported (*19*). In contrast, even though neuronal outgrowth has been explored in ALS (*20*), our identification of genetic risk underlying *neuronal morphogenesis* was novel. Indeed, the combination of *membrane trafficking* and *neuronal morphogenesis* may be a driving force of the disease pathogenesis. The defining feature of motor neurons is the length of their axons, projections that require specialized long-range transport and efficient cytoskeletal dynamics for the maintenance of synaptic connections (*21*). Similarly, *signal transduction mediated by ribonucleotides* is a broad term encompassing ion channel transport that regulates signal transmission at synapses. Disruption of this process leads to hyperexcitability, a phenomenon that has been observed in ALS patients (*22*). We speculate that broadly expressed genes lead to selective damage due to the high reliance of motor neurons on cellular transport, morphogenesis, and axonal ion channels compared to other cell types.

Our data did not detect biological pathways that have been previously implicated in the pathogenesis of familial ALS, such as *nucleocytoplasmic transport (23)* and *excitotoxicity* (*24*). These cellular processes may only operate in specific genetic forms of ALS, such as *C9orf72*-related or *SOD1*-related cases. A more likely explanation is that rare and low-frequency variants not captured by our methodology, significantly contribute to those pathways. For this reason, we cannot rule these biological processes out as relevant to the pathogenesis of ALS. Future analyses of more substantial datasets that include whole-genome sequencing data may implicate them.

One of our study’s strengths is that we were able to distinguish differential pathways operating in *C9orf72* expansion carriers versus non-carriers. Interestingly, the *autophagosome* pathway was only significant in the analysis of the *C9orf72* expansion carriers. The *C9orf72* protein is a known regulator of autophagy; hence it is not surprising that a higher burden of ALS genetic risk was found within autophagy genes in *C9orf72* expansion carriers versus non-carriers. This is the first time that autophagy-related processes have been implicated in *C9orf72* biology from a genetic perspective. The hexanucleotide repeat expansion is known to influence the expression of the *C9orf72* gene, irrespective of reported biology involving dipeptide repeats and toxic RNA species arising directly from the repeat expansion (*25*), which reinforces the importance of our findings. The C9orf72 protein was also recently found to play a role in neuronal and dendritic morphogenesis in ALS through the promotion of autophagy (*26*).

Our rigorous approach using multiple human and mouse transcriptome datasets identified *cortical GABAergic interneurons* and *oligodendrocytes* as the cell types central to ALS. These findings are consistent with published literature. For example, alteration in inhibitory signaling through GABAergic interneurons contributes to neural hyperexcitability, an early event in ALS pathogenesis (*22*). Oligodendrocytes from sporadic and familial *SOD1* ALS exert a harmful effect on motor neurons through the secretion of toxic factors (*27*). Even though these cell types were previously linked with toxicity in ALS, our study indicates that oligodendrocytes incorporate a significant proportion of ALS genetic risk. This initial finding supports the idea that these cells directly contribute to the disease pathogenesis, rather than merely playing a secondary role in the disease progression.

Our results show the power of data-driven approaches to nominate aspects of the nervous system for additional scrutiny. Nevertheless, our study has limitations. Though we analyzed 78,500 individuals in the current study, our power to detect pathways remains limited. This lack of power primarily stems from the genetic architecture of ALS, which is known to conform to the rare disease-rare variant paradigm (*28*). By design, pathway analysis focuses on common variants with a frequency greater than 1%, but we know that the contribution of common variants to ALS risk is modest (*6*). Furthermore, our approach is based on intragenic variants, even though intergenic mutations can affect gene expression. We have overcome this power limitation by performing multiple rounds of replication in both the pathway analysis and cell type analysis to ensure accuracy and validity. The detected pathways and cell types represent potent aspects of the ALS disease process, but additional critical cellular mechanisms will undoubtedly be found using more extensive datasets.

Another limitation is the Molecular Signatures Database that we used to define the pathways in our analysis. This collection is incomplete, especially for neuronal and glial pathways (*29*). As our understanding evolves and more single-cell expression datasets become available, it may be worthwhile to reevaluate our GWAS data periodically. To facilitate this, we have made the programming code needed to perform the analysis publicly available. We also created an interactive, online resource that enables the research community to explore the contribution of pathways and cell types to ALS risk (https://lng-nia.shinyapps.io/ALS-Pathways/).

In conclusion, we demonstrate the utility of data-driven approaches to dissect the molecular basis of complex diseases such as ALS. Our stringent approach points to *neuron projection morphogenesis, membrane trafficking*, and *signal transduction mediated by ribonucleotides* as primary drivers of motor neuron degeneration in ALS. It also nominates *cortical GABAergic interneurons* and *oligodendrocytes* as central to the pathogenesis of this fatal neurological disease.

## MATERIAL AND METHODS

### EXPERIMENTAL DESIGN

#### Study design

We used a three-stage study design to identify pathways relevant to ALS risk (see **Fig. 1** for workflow). To ensure accuracy, we compiled the available ALS genomic data into three independent datasets for analyses. The reference dataset consisted of summary statistics from a previous published GWAS involving 12,577 cases and 23,475 controls of European ancestry (publicly available from databrowser.projectmine.com)(*6*). We used the summary statistics from this reference dataset to define risk allele weights for constructing polygenic risk scores within pathways defined by the Molecular Signatures Database.

The remaining data consisted of individual-level genotype and phenotype data from 8,073 ALS cases and 35,184 control subjects of European ancestry that we recently published (*7*). We randomly split these data in a 70%-to-30% ratio into a “training dataset” containing 5,653 cases and 24,629 control subjects, and a “replication dataset” consisting of 2,420 patients and 10,555 controls subjects. We used the regression model generated from the reference data to construct and test polygenic risk scores within the training data. The replication dataset was used to validate our training data findings. There was no sample overlap between the reference, training, or replication datasets.

#### Human subjects

All patients included in our analysis had been diagnosed with ALS according to the El Escorial criteria by a neurologist specializing in ALS. The demographics of the cohorts are listed in supplementary table 3. Written consent was obtained from all individuals enrolled in this study, and the study was approved by the institutional review board of the National Institute on Aging (protocol number 03-AG-N329).

The human samples for single-nucleus RNA-seq consisted of frozen frontal cortex post-mortem samples obtained from 16 neurologically healthy donors. The subjects were between 16 and 61 years of age (median age = 36, male: female ratio = 1:1). The samples were acquired from the University of Maryland Brain and Tissue Bank through the NIH NeuroBioBank.

### METHOD DETAILS

#### Gene set selection and pathway analysis

The Molecular Signatures Database (MSigDB database v6.2, http://software.broadinstitute.org/gsea/downloads_archive.jsp) is a compilation of annotated gene sets designed for gene enrichment and pathway analysis. This database is divided into eight collections (*30, 31*), and we focused our efforts on three of these compilations that have been validated previously (*8*)(*9*): (1) Hallmark gene sets representing well-defined biological processes (n = 50); (2) Curated gene sets representing pathways annotated by various sources such as online pathway databases, the biomedical literature, and manual curation by domain experts (n = 1,330); and (3) Gene ontology gene sets consisting of pathways annotated with the same gene ontology term (n = 5,967). The last collection is subdivided into biological processes, cellular components, and molecular functions (see ***SI Appendix*, Fig. S1**).

#### Quality control of reference and target datasets

The target dataset consisted of individual-level genotype and phenotype data in the PLINK binary file format. Only variants with an imputation quality (R_2_) greater than 0.8 were included in the analysis. To ensure that the *C9orf72* gene was correctly represented in the dataset, we removed 120 kb upstream and downstream of *C9orf27*, and we replaced rs3849943 (located outside *C9orf72*) with rs2453555 (located within intron 3). After these filters, 5,421,177 variants remained in the training dataset. From these, we selected 268,431 variants with an association p-value in the reference dataset less than or equal to 0.05. Next, we applied the default clumping parameters outlined in the *PRSice2* software package (*32*) (version 2.1.1, R_2_ = 0.1, and a 250 kb window). This clumping process yielded 27,176 variants that were then used for polygenic risk score analysis.

#### Polygenic risk score generation

Polygenic risk scores were calculated based on the weighted allele dosages as implemented in *PRSIce2* using the no-clump flag. A key advantage of this approach is that it allows variants below the typical GWAS significance threshold of 5.0×10_-8_ to be included in the analysis. For the training dataset, 1,000 permutations were used to generate empirical p-value estimates for each GWAS-derived p-value. Each permutation test in the training dataset provided a Nagelkerke’s pseudo R_2_ value after adjusting for an estimated ALS prevalence of 5 per 100,000 of the population (*33*). Sex, age at onset, and eigenvectors one to twenty were included as covariates in the model.

To test the contribution of known ALS GWAS genetic risk loci to our pathways, we included the following risk variants as covariates in the replication testing: rs10463311 (*TNIP1*), rs2453555 (*C9orf72*), rs113247976 (*KIF5A*), rs74654358 (*TBK1*), and rs12973192 (*UNC13A*). The variant rs75087725 corresponding to the *C21orf2* gene was not included as this variant has a low imputation quality score (R_2_ < 0.8). Also, although rs12973192 is the variant nominated as the *UNC13A* GWAS hit, it was replaced by the clumping algorithm in favor of rs7849703.

Polygenic risk scores were then tested in the replication phase using the --score command implemented in PLINK v1.9 (*34*). Polygenic risk scores were calculated, incorporating the risk variants from the pathways nominated in the discovery phase. Risk allele dosages were counted (giving a dose of two if homozygous for the risk allele, one if heterozygous, and zero if homozygous for the reference allele). All SNPs were weighted by the log-odds ratios obtained from the reference dataset with a greater weight given to alleles with higher risk estimates. Polygenic risk scores were converted to Z scores for easier interpretation. Logistic regressions were performed to evaluate the association between the pathway-specific polygenic risk score of interest with ALS as the outcome. Gene sets/pathways containing less than 20 SNPs were discarded.

An example of the polygenic risk score procedure is as follows: the Molecular Signature Database lists 79 genes as part of the *autophagosome* pathway. After applying our filtering methods, 50 variants were located within these genes that achieved a p-value of less than 0.05 in the reference GWAS. These 50 variants were used to calculate the polygenic risk score of the *autophagosome* pathway in the training dataset. We scaled the risk allele dosages of these variants using the beta estimates obtained from the reference dataset. Finally, we evaluated these 50 variants in the independent replication dataset.

#### Semantic similarity analysis of gene ontology terms

The *GoSemSim* function from the *GoSemSim* R package (version 2.8.0) was used to calculate the semantic similarity between sets of gene ontology terms (*35*). This algorithm applies Wang’s method based on a graph-based strategy using the topology of the gene ontology graph structure. Hierarchical clustering based on similarity scores was performed to separate groups of gene ontology terms, and the groups were labeled using a representative term. To obtain the representative term, we used the function *tree map* from REVIGO (http://revigo.irb.hr) (*11*). Additionally, SNPs from the cellular component significant set and the molecular function significant set were further subjected to enrichment analysis to dissect biological function. The function *g:GOSt* from *g:ProfileR* (Raudvere et al., 2019) (https://biit.cs.ut.ee/gprofiler/gost) was used to detect the top three REACTOME enriched pathways (***SI Appendix*, Fig S2)**.

#### Mendelian randomization analysis

To identify genes within the thirteen significant pathways that drive the risk of ALS, we exploited the known tendency of SNPs associated with disease also to be associated with gene eQTL (*36*). We applied summary-data-based Mendelian randomization as implemented in the *SMR* software package (http://cnsgenomics.com/software/smr) (*37*) to the genes within the thirteen significant pathways. This approach used estimates for cis-eQTLs obtained from a sizeable eQTL meta-analysis performed in blood (*13*) and brain (*14*). Brain expression datasets include estimates for cis-expression from the Genotype-Tissue Expression (GTEx) Consortium (v6; whole blood and ten brain regions) (*15*), the Common Mind Consortium (dorsolateral prefrontal cortex) (*16*) and the Religious Orders Study and Memory and Aging Project (ROSMAP) (*17*). This methodology used summary-level data from GWAS and eQTL studies to test for pleiotropic association. Wald ratios were generated for each instrumental variable SNP tagging a cis-eQTL (defined as probes within a gene that met an eQTL p-value of at least 5×10_-8_ in the original study). Linkage pruning and clumping were carried out using default *SMR* protocols. The p-values per instrument substrate were adjusted by FDR. SNPs with a HEIDI p-value of less than 0.01 were excluded on the grounds of pleiotropy.

#### Nuclei isolation

Approximately 100mg of tissue was homogenized in cold Lysis Buffer (Nuclei PURE Lysis Buffer/1mM DTT/0.1% Triton X-100, Sigma Aldrich) in a Dounce homogenizer. The homogenate was transferred to a 50mL conical tube, vortexed for 2-3 seconds, and incubated for 10 minutes on ice in a total of 10mL Lysis Buffer. Tissue lysate was resuspended with 18mL of cold 1.8M Sucrose Cushion Solution and layered slowly over 10mL of cold 1.8M Sucrose Cushion Solution (Sigma Aldrich) in an ultracentrifuge tube (Beckman Coulter) on ice. Samples were centrifuged for 45 minutes at 30,000*g* at 4 °C. Pelleted nuclei were resuspended in 1mL cold Nuclei Suspension Buffer (NSB, PBS/0.01%, BSA/0.1% (New England BioLabs), and SUPERase-In RNase inhibitor (ThermoFisher Scientific) (*18*). The nuclei suspension was mixed with an additional 4mL of cold NSB and centrifuged for 5 minutes at 500*g* at 4 °C. After a second wash in 5mL cold NSB, nuclei were resuspended in 100-200μL of cold NSB and counted on an automated cell counter (Bio-Rad). The concentration of the nuclei suspension was adjusted to ∼1000 nuclei/μL.

#### Single-nucleus RNA sequencing

The extracted nuclei were submitted to the Single Cell Analysis Facility (Center for Cancer Research, National Cancer Institute) for single-cell RNA sequencing. Sequencing libraries were constructed using the Chromium Single Cell Gene Expression Solution v3 (10x Genomics). The libraries were pooled and loaded at a concentration of 1.8pM with 10% PhiX spike-in for sequencing on an Illumina NextSeq 550 System using Illumina NextSeq 150 Cycle Hi-Output v2.5 kits (Illumina) to achieve a targeted read depth of ∼33,000 reads per nucleus. The resulting FASTQ files were aligned and counted using *Cell Ranger* software v3 (10x Genomics), generating feature-barcode matrices. One donor was sequenced in triplicate, and two donors were sequenced in duplicate to produce a total of 21 single-cell RNA-seq datasets.

These datasets were normalized using *SCTransform* v0.2.1 (*38*) and integrated by pair-wise comparison of anchor gene expression (*39*) within the *Seurat* package v3.1 (*40*) in R. Shared-nearest-neighbor-based clustering was used to identify distinct cell clusters, which were then manually assigned cell type identities based on differential expression of known cell type marker genes (*41*)(*42*).

#### Cell type-specific risk

We used in-house-generated single-nuclei RNA-seq data obtained from the human frontal cortex. These data were based on 161,225 nuclei transcriptomes from 16 neurologically healthy donors. We calculated the specificity of expression for each gene within each cell type, following a previously published methodology (*10*). These values range from zero to one and represent the proportion of the total expression of a gene found in one cell type compared to all cell types. For example, if a gene has a score of one in a particular cell type, it means that it is only expressed in this cell type. If a gene has a score of zero in a given cell type, it means that it is not expressed in that cell type (*10*).

Overall, we assessed the variance explained by the polygenic risk score (pseudo-R_2_) in thirty-four human brain cell types. We obtained the pseudo-R_2_ value using the merged training and replication datasets (8,016 cases and 34,432 controls). Next, we applied a linear regression model to evaluate if more of the variance explained by polygenic risk score is attributable to the genes that were more specific to each cell type (p-value of the hypothesis test in the model < 0.05).

For replication, we used publicly available droplet single-nucleus RNA-seq (DroNc-seq) data consisting of 19,550 nuclei obtained from four frozen, post-mortem samples of human hippocampus and three samples from prefrontal cortex (*43*). The specificity matrix for this dataset was obtained from https://github.com/RHReynolds/MarkerGenes/tree/master/specificity_matrices.

### STATISTICAL ANALYSIS

All statistics were performed using R and Plink version 1.9. Polygenic risk scores were calculated using the *PRSIce2* algorithm, and 1,000 permutations were used to generate empirical p-value estimates as described in the Methods. A linear regression model was used in the cell type analysis to evaluate the variance explained by the polygenic risk score attributable to genes specific to each cell type. Significance thresholds were set at p < 0.05 (FDR-corrected in the training dataset and raw p-value in the replication dataset). Power calculations were performed by estimating the variance in the training dataset using the *estimatePolygenicModel* function within the *AVENGEME* v1 package (https://github.com/DudbridgeLab/avengeme/)(*44*), and then determining the power of the polygenic risk score to predict disease status in the replication dataset using the *polygenescore* function. Data statistics are detailed in figure legends, and statistical values are listed in the Results section.

## DATA AND CODE AVAILABILITY

The results are available online at https://lng-nia.shinyapps.io/ALS-Pathways/. This interactive web portal includes data for the 7,347 pathways and gene sets of the Molecular Signatures Database, as well as data for twenty-four cell types. The programming code and genetic data used for this study are available at https://github.com/sarasaezALS/ALS-Pathway and dbGaP (https://www.ncbi.nlm.nih.gov/projects/gap/cgi-bin/study.cgi?study_id=phs000101.v5.p1).

## ACKNOWLEDGEMENTS

This work was supported in part by the Intramural Research Program of the NIH, National Institute on Aging (Z01-AG000949-02), by the National Institute of Neurological Disorders and Stroke, and by Merck Sharp & Dohme Corp., a subsidiary of Merck & Co., Inc., Kenilworth, NJ, USA. Bryan J. Traynor received additional support from the Center for Disease Control and Prevention, the Muscular Dystrophy Association, Microsoft Research, the Packard Center for ALS Research at Johns Hopkins, and the ALS Association. This study utilized the high-performance computational capabilities of the Biowulf Linux cluster at the NIH (http://hpc.nih.gov). The authors would like to thank the Project MinE GWAS Consortium. This study also used genotype and clinical data from the Wellcome Trust Case Control Consortium, and the HyperGenes Consortium. We thank the patients and research subjects who contributed samples for this study.

This work was supported in part by the Italian Ministry of Health (Ministero della Salute, Ricerca Sanitaria Finalizzata, grant RF-2016-02362405); the European Commission’s Health Seventh Framework Programme (FP7/2007-2013 under grant agreement 259867); the Italian Ministry of Education, University and Research (Progetti di Ricerca di Rilevante Interesse Nazionale, PRIN, grant 2017SNW5MB); the Joint Programme - Neurodegenerative Disease Research (Brain-Mend project) granted by Italian Ministry of Education, University and Research, and the Canadian Consortium on Neurodegeneration in Aging (CCNA). This study was performed under the Department of Excellence grant of the Italian Ministry of Education, University and Research to the ‘Rita Levi Montalcini’ Department of Neuroscience, University of Torino, Italy.

This study used DNA samples and clinical data from the Target ALS Human Postmortem Tissue Core, the NINDS Repository at Coriell, the North East ALS (NEALS) Consortium Biorepository, the New York Brain Bank-The Taub Institute, Columbia University, Department of Veterans Affairs Biorepository Brain Bank (grant #BX002466; C.B.B.), the Baltimore Longitudinal Study of Aging (BLSA), and the Johns Hopkins University Alzheimer’s Disease Research Center, the NICHD Brain and Tissue Bank for Developmental Disorders at the University of Maryland.

The InCHIANTI study baseline (1998–2000) was supported as a ‘‘targeted project’’ (ICS110.1/RF97.71) by the Italian Ministry of Health and in part by the United States National Institute on Aging (contracts 263 MD 9164 and 263 MD 821336). The InCHIANTI follow-up 1 (2001–2003) was funded by the United States National Institute on Aging (contracts N.1-AG-1-1 and N.1-AG-1-2111), and the InCHIANTI follow-ups 2 and 3 studies (2004–2010) were financed by the United States National Institute on Aging (contract N01-AG-5-0002). The dataset(s) used for the analyses described in this manuscript were obtained from the Age-Related Eye Disease Study (AREDS) Database found at https://www.nei.nih.gov/research/clinical-trials/age-related-eye-disease-study-aredsthrough dbGaP accession number phs000001.v3.p1. Funding support for AREDS was provided by the National Eye Institute (N01-EY-0-2127). We would like to thank the AREDS participants and the AREDS Research Group for their valuable contribution to this research. The Framingham Heart Study is conducted and supported by the National Heart, Lung, and Blood Institute (NHLBI) in collaboration with Boston University (Contract No. N01-HC-25195 and HHSN268201500001I). This manuscript was not prepared in collaboration with investigators of the Framingham Heart Study and does not necessarily reflect the opinions or views of the Framingham Heart Study, Boston University, or NHLBI. Funding to support the Omni cohort recruitment, retention and examination was provided by NHLBI Contract N01-HC-25195 and HHSN268201500001I, as well as NHLBI grants R01-HL070100, R01-HL076784, R01-HL-49869, and U01-HL-053941. Research support to collect data and develop an application to support this project was provided by 3P50CA093459, 5P50CA097007, 5R01ES011740, and 5R01CA133996.

The WHI program is funded by the National Heart, Lung, and Blood Institute, National Institutes of Health, U.S. Department of Health and Human Services through contracts HHSN268201600018C, HHSN268201600001C, HHSN268201600002C, HHSN268201600003C, and HHSN268201600004C. This manuscript was not prepared in collaboration with investigators of the WHI, has not been reviewed and/or approved by the Women’s Health Initiative (WHI), and does not necessarily reflect the opinions of the WHI investigators or the NHLBI. Funding support for WHI GARNET was provided through the NHGRI Genomics and Randomized Trials Network (GARNET) (Grant Number U01 HG005152). Assistance with phenotype harmonization and genotype cleaning, as well as with general study coordination, was provided by the GARNET Coordinating Center (U01 HG005157). Assistance with data cleaning was provided by the National Center for Biotechnology Information. Funding support for genotyping, which was performed at the Broad Institute of MIT and Harvard, was provided by the NIH Genes, Environment and Health Initiative [GEI] (U01 HG004424). The datasets used for the analyses described in this manuscript were obtained from dbGaP at http://www.ncbi.nlm.nih.gov/sites/entrez?db=gap through dbGaP accession phs000001, phs000007, phs000187, phs000196, phs000200, phs000315, phs000675, phs000248, phs000292, phs000304, phs000368, phs000372, phs000394, phs000397, phs000404, phs000421, phs000428, phs000454, phs000615, phs000801, and phs000869.

The authors acknowledge the contribution of data from Hepatitis C Pathogenesis and the Human Genome supported by 1×01HG005271-01 and R01DA013324 and accessed through dbGAP to the analysis presented in this publication.

Funding support for the Genes and Blood Clotting Study was provided through the NIH/NHLBI (R37 HL039693). The Genes and Blood Clotting Study is one of the Phase 3 studies as part of the Gene Environment Association Studies (GENEVA) under GEI. Assistance with genotype cleaning was provided by the GENEVA Coordinating Center (U01 HG004446). Funding support for DNA extraction and genotyping, which was performed at the Broad Institute, was provided by NIH/NHLBI (R37 HL039693). Additional support was provided by the Howard Hughes Medical Institute.

The dataset(s) used for the analyses described in this manuscript were obtained from the database of Genotype and Phenotype (dbGaP) found at http://www.ncbi.nlm.nih.gov/gap through dbGaP accession number phs000368. Samples and associated phenotype data for the Genome-Wide Association Scan [GWAS] of Polycystic Ovary Syndrome Phenotypes were provided by Andrea Dunaif, M.D.

The authors acknowledge the contribution of data from Genetic Architecture of Smoking and Smoking Cessation accessed through dbGAP. Funding support for genotyping, which was performed at the Center for Inherited Disease Research (CIDR), was provided by 1 X01 HG005274-01. CIDR is fully funded through a federal contract from the National Institutes of Health to The Johns Hopkins University, contract number HHSN268200782096C. Assistance with genotype cleaning, as well as with general study coordination, was provided by the Gene Environment Association Studies (GENEVA) Coordinating Center (U01 HG004446). Funding support for collection of datasets and samples was provided by the Collaborative Genetic Study of Nicotine Dependence (COGEND; P01 CA089392) and the University of Wisconsin Transdisciplinary Tobacco Use Research Center (P50 DA019706, P50 CA084724).

The dataset(s) used for the analyses described in this manuscript were obtained from the Genetics of Fuchs’ Endothelial Corneal Dystrophy (FECD) Study through dbGaP accession number phs000421. The grants that have funded the enrollment of the cases and controls to be used in this GWAS are:R01EY016514 (DUEC, PI: Gordon Klintworth), R01EY016482 (CWRU, PI: Sudha Iyengar), and 1×01HG006619-01 (PI: Sudha Iyengar, Natalie Afshari). We would like to thank the FECD participants and the FECD Research Group for their valuable contribution to this research The authors acknowledge the contribution of data from CIDR-NIDA Study of HIV Host Genetics accessed through dbGAP. Funding support for genotyping, which was performed at the Center for Inherited Disease Research (CIDR), was provided by 1 X01 HG005275-01A1. CIDR is fully funded through a federal contract from the National Institutes of Health to The Johns Hopkins University, contract number HHSN268200782096C. Funding support for collection of datasets and samples was provided by NIDA grants R01DA026141(Johnson); R01DA004212 (Watters); U01DA006908 (Watters); R01DA009532 (Bluthenthal); as well as the San Francisco Department of Public Health; SAMHSA; and HRSA.

The Genome-Wide Association Study (GWAS) of Non-Hodgkin Lymphoma (NHL) project was supported by the intramural program of the Division of Cancer Epidemiology and Genetics (DCEG), National Cancer Institute (NCI), National Institutes of Health (NIH). The datasets have been accessed through the NIH database for Genotypes and Phenotypes (dbGaP) under accession # phs000801. A full list of acknowledgements can be found in the supplementary note (Berndt SI et al., Nature Genet., 2013, PMID: 23770605).

This study made use of data generated by investigators in the BEACON consortium through a grant funded by the US National Institutes of Health (NIH) (RO1CA136725) to Thomas L. Vaughan and David C. Whiteman (multiple PIs). In support of this work, T.L.V. was also supported by NIH grant KO5CA124911 and D.C.W. by a Future Fellowship grant FT0990987 from the Australia Research Council. Additional collaborators, sources of support and origin of the data and biospecimens are listed in the following publication: Levine DM, Ek WE, Zhang R, Liu X, Onstad L, Sather C, et al. A genome-wide association study identifies new susceptibility loci for esophageal adenocarcinoma and Barrett’s esophagus. Nat Genet. 2013 Dec;45(12):1487– 93.

## Author contributions

Conception and design of the study: SSA, SBC, MAN, and BJT.

Acquisition and analysis of data: SSA, SBC, RGL, MAN, RC, SWC, AC, and BJT.

Designed and implemented the online portal: KJ

Drafted the manuscript: SSA and BJT.

Edit the manuscript: all the other authors commented on and edited the manuscript.

## References

1. K. C. Arthur, A. Calvo, T. R. Price, J. T. Geiger, A. Chiò, B. J. Traynor, Projected increase in amyotrophic lateral sclerosis from 2015 to 2040. Nat. Commun. 7, 12408 (2016).

2. R. H. Brown, A. Al-Chalabi, Amyotrophic Lateral Sclerosis. N. Engl. J. Med. 377, 162–172 (2017).

3. D. B. Medinas, P. Rozas, F. Martínez Traub, U. Woehlbier, R. H. Brown, D. A. Bosco, C. Hetz, Endoplasmic reticulum stress leads to accumulation of wild-type SOD1 aggregates associated with sporadic amyotrophic lateral sclerosis. Proc. Natl. Acad. Sci. U. S. A. 115, 8209–8214 (2018).

4. C.-C. Chou, Y. Zhang, M. E. Umoh, S. W. Vaughan, I. Lorenzini, F. Liu, M. Sayegh, P. G. Donlin-Asp, Y. H. Chen, D. M. Duong, N. T. Seyfried, M. A. Powers, T. Kukar, C. M. Hales, M. Gearing, N. J. Cairns, K. B. Boylan, D. W. Dickson, R. Rademakers, Y.-J. Zhang, L. Petrucelli, R. Sattler, D. C. Zarnescu, J. D. Glass, W. Rossoll, TDP-43 pathology disrupts nuclear pore complexes and nucleocytoplasmic transport in ALS/FTD. Nat. Neurosci. 21, 228–239 (2018).

5. V. Valenzuela, M. Nassif, C. Hetz, Unraveling the role of motoneuron autophagy in ALS. Autophagy. 14, 733–737 (2018).

6. W. van Rheenen, A. Shatunov, A. M. Dekker, R. L. McLaughlin, F. P. Diekstra, S. L. Pulit, R. A. A. van der Spek, U. Võsa, S. de Jong, M. R. Robinson, J. Yang, I. Fogh, P. T. van Doormaal, G. H. P. Tazelaar, M. Koppers, A. M. Blokhuis, W. Sproviero, A. R. Jones, K. P. Kenna, K. R. van Eijk, O. Harschnitz, R. D. Schellevis, W. J. Brands, J. Medic, A. Menelaou, A. Vajda, N. Ticozzi, K. Lin, B. Rogelj, K. Vrabec, M. Ravnik-Glavač, B. Koritnik, J. Zidar, L. Leonardis, L. D. Grošelj, S. Millecamps, F. Salachas, V. Meininger, M. de Carvalho, S. Pinto, J. S. Mora, R. Rojas-García, M. Polak, S. Chandran, S. Colville, R. Swingler, K. E. Morrison, P. J. Shaw, J. Hardy, R. W. Orrell, A. Pittman, K. Sidle, P. Fratta, A. Malaspina, S. Topp, S. Petri, S. Abdulla, C. Drepper, M. Sendtner, T. Meyer, R. A. Ophoff, K. A. Staats, M. Wiedau-Pazos, C. Lomen-Hoerth, V. M. Van Deerlin, J. Q. Trojanowski, L. Elman, L. McCluskey, A. N. Basak, C. Tunca, H. Hamzeiy, Y. Parman, T. Meitinger, P. Lichtner, M. Radivojkov-Blagojevic, C. R. Andres, C. Maurel, G. Bensimon, B. Landwehrmeyer, A. Brice, C. A. M. Payan, S. Saker-Delye, A. Dürr, N. W. Wood, L. Tittmann, W. Lieb, A. Franke, M. Rietschel, S. Cichon, M. M. Nöthen, P. Amouyel, C. Tzourio, J.-F. Dartigues, A. G. Uitterlinden, F. Rivadeneira, K. Estrada, A. Hofman, C. Curtis, H. M. Blauw, A. J. van der Kooi, M. de Visser, A. Goris, M. Weber, C. E. Shaw, B. N. Smith, O. Pansarasa, C. Cereda, R. Del Bo, G. P. Comi, S. D’Alfonso, C. Bertolin, G. Sorarù, L. Mazzini, V. Pensato, C. Gellera, C. Tiloca, A. Ratti, A. Calvo, C. Moglia, M. Brunetti, S. Arcuti, R. Capozzo, C. Zecca, C. Lunetta, S. Penco, N. Riva, A. Padovani, M. Filosto, B. Muller, R. J. Stuit, PARALS Registry, SLALOM Group, SLAP Registry, FALS Sequencing Consortium, SLAGEN Consortium, NNIPPS Study Group, I. Blair, K. Zhang, E. P. McCann, J. A. Fifita, G. A. Nicholson, D. B. Rowe, R. Pamphlett, M. C. Kiernan, J. Grosskreutz, O. W. Witte, T. Ringer, T. Prell, B. Stubendorff, I. Kurth, C. A. Hübner, P. N. Leigh, F. Casale, A. Chio, E. Beghi, E. Pupillo, R. Tortelli, G. Logroscino, J. Powell, A. C. Ludolph, J. H. Weishaupt, W. Robberecht, P. Van Damme, L. Franke, T. H. Pers, R. H. Brown, J. D. Glass, J. E. Landers, O. Hardiman, P. M. Andersen, P. Corcia, P. Vourc’h, V. Silani, N. R. Wray, P. M. Visscher, P. I. W. de Bakker, M. A. van Es, R. J. Pasterkamp, C. M. Lewis, G. Breen, A. Al-Chalabi, H. van den Berg, J. H. Veldink, Genome-wide association analyses identify new risk variants and the genetic architecture of amyotrophic lateral sclerosis. Nat. Genet. 48, 1043–1048 (2016).

7. A. Nicolas, K. P. Kenna, A. E. Renton, N. Ticozzi, F. Faghri, R. Chia, J. A. Dominov, B. J. Kenna, A. Nalls, P. Keagle, A. M. Rivera, W. van Rheenen, N. A. Murphy, J. J. F. A. van Vugt, J. T. Geiger, R. A. Van der Spek, H. A. Pliner, Shankaracharya, B. N. Smith, G. Marangi, S. D. Topp, Y. Abramzon, A. S. Gkazi, J. D. Eicher, A. Kenna, ITALSGEN Consortium, G. Mora, A. Calvo, L. Mazzini, N. Riva, J. Mandrioli, C. Caponnetto, S. Battistini, P. Volanti, V. La Bella, F. L. Conforti, G. Borghero, S. Messina, I. L. Simone, F. Trojsi, F. Salvi, F. O. Logullo, S. D’Alfonso, L. Corrado, M. Capasso, L. Ferrucci, Genomic Translation for ALS Care (GTAC) Consortium, C. de A. M. Moreno, S. Kamalakaran, D. B. Goldstein, ALS Sequencing Consortium, A. D. Gitler, T. Harris, R. M. Myers, NYGC ALS Consortium, H. Phatnani, R. L. Musunuri, U. S. Evani, A. Abhyankar, M. C. Zody, Answer ALS Foundation, J. Kaye, S. Finkbeiner, S. K. Wyman, A. LeNail, L. Lima, E. Fraenkel, C. N. Svendsen, L. M. Thompson, J. E. Van Eyk, J. D. Berry, T. M. Miller, S. J. Kolb, M. Cudkowicz, E. Baxi, Clinical Research in ALS and Related Disorders for Therapeutic Development (CReATe) Consortium, M. Benatar, J. P. Taylor, E. Rampersaud, G. Wu, J. Wuu, SLAGEN Consortium, G. Lauria, F. Verde, I. Fogh, C. Tiloca, G. P. Comi, G. Sorarù, C. Cereda, French ALS Consortium, P. Corcia, H. Laaksovirta, L. Myllykangas, L. Jansson, M. Valori, J. Ealing, H. Hamdalla, S. Rollinson, S. Pickering-Brown, R. W. Orrell, K. C. Sidle, A. Malaspina, J. Hardy, A. B. Singleton, J. O. Johnson, S. Arepalli, P. C. Sapp, D. McKenna-Yasek, M. Polak, S. Asress, S. Al-Sarraj, A. King, C. Troakes, C. Vance, J. de Belleroche, F. Baas, A. L. M. A. Ten Asbroek, J. L. Muñoz-Blanco, D. G. Hernandez, J. Ding, J. R. Gibbs, S. W. Scholz, M. K. Floeter, R. H. Campbell, F. Landi, R. Bowser, S. M. Pulst, J. M. Ravits, D. J. L. MacGowan, J. Kirby, E. P. Pioro, R. Pamphlett, J. Broach, G. Gerhard, T. L. Dunckley, C. B. Brady, N. W. Kowall, J. C. Troncoso, I. Le Ber, K. Mouzat, S. Lumbroso, T. D. Heiman-Patterson, F. Kamel, L. Van Den Bosch, R. H. Baloh, T. M. Strom, T. Meitinger, A. Shatunov, K. R. Van Eijk, M. de Carvalho, M. Kooyman, B. Middelkoop, M. Moisse, R. L. McLaughlin, M. A. Van Es, M. Weber, K. B. Boylan, M. Van Blitterswijk, R. Rademakers, K. E. Morrison, A. N. Basak, J. S. Mora, V. E. Drory, P. J. Shaw, M. R. Turner, K. Talbot, O. Hardiman, K. L. Williams, J. A. Fifita, G. A. Nicholson, I. P. Blair, G. A. Rouleau, J. Esteban-Pérez, A. García-Redondo, A. Al-Chalabi, Project MinE ALS Sequencing Consortium, E. Rogaeva, L. Zinman, L. W. Ostrow, N. J. Maragakis, J. D. Rothstein, Z. Simmons, J. Cooper-Knock, A. Brice, S. A. Goutman, E. L. Feldman, S. B. Gibson, F. Taroni, A. Ratti, C. Gellera, P. Van Damme, W. Robberecht, P. Fratta, M. Sabatelli, C. Lunetta, A. C. Ludolph, P. M. Andersen, J. H. Weishaupt, W. Camu, J. Q. Trojanowski, V. M. Van Deerlin, R. H. Brown Jr, L. H. van den Berg, J. H. Veldink, M. B. Harms, J. D. Glass, D. J. Stone, P. Tienari, V. Silani, A. Chiò, C. E. Shaw, B. J. Traynor, J. E. Landers, Genome-wide Analyses Identify KIF5A as a Novel ALS Gene. Neuron. 97, 1268–1283.e6 (2018).

8. A. Liberzon, C. Birger, H. Thorvaldsdóttir, M. Ghandi, J. P. Mesirov, P. Tamayo, The Molecular Signatures Database Hallmark Gene Set Collection. Cell Systems. 1 (2015), pp. 417–425.

9. J. Bubier, D. Hill, G. Mukherjee, T. Reynolds, E. J. Baker, A. Berger, J. Emerson, J. A. Blake, E. J. Chesler, Curating gene sets: challenges and opportunities for integrative analysis. Database. 2019 (2019), doi: 10.1093/database/baz036.

10. N. G. Skene, J. Bryois, T. E. Bakken, G. Breen, J. J. Crowley, H. A. Gaspar, P. Giusti-Rodriguez, R. D. Hodge, J. A. Miller, A. B. Muñoz-Manchado, M. C. O’Donovan, M. J. Owen, A. F. Pardiñas, J. Ryge, J. T. R. Walters, S. Linnarsson, E. S. Lein, Major Depressive Disorder Working Group of the Psychiatric Genomics Consortium, P. F. Sullivan, J. Hjerling-Leffler, Genetic identification of brain cell types underlying schizophrenia. Nat. Genet. 50, 825–833 (2018).

11. F. Supek, M. Bošnjak, N. Škunca, T. Šmuc, REVIGO summarizes and visualizes long lists of gene ontology terms. PLoS One. 6, e21800 (2011).

12. D. L. Nicolae, E. Gamazon, W. Zhang, S. Duan, M. Eileen Dolan, N. J. Cox, Trait-Associated SNPs Are More Likely to Be eQTLs: Annotation to Enhance Discovery from GWAS. PLoS Genet. 6, e1000888 (2010).

13. U. Võsa, A. Claringbould, H.-J. Westra, M. J. Bonder, P. Deelen, B. Zeng, H. Kirsten, A. Saha, R. Kreuzhuber, S. Kasela, N. Pervjakova, I. Alvaes, M.-J. Fave, M. Agbessi, M. Christiansen, R. Jansen, I. Seppälä, L. Tong, A. Teumer, K. Schramm, G. Hemani, J. Verlouw, H. Yaghootkar, R. Sönmez, A. Brown, V. Kukushkina, A. Kalnapenkis, S. Rüeger, E. Porcu, J. Kronberg-Guzman, J. Kettunen, J. Powell, B. Lee, F. Zhang, W. Arindrarto, F. Beutner, BIOS Consortium, H. Brugge, i2QTL Consortium, J. Dmitreva, M. Elansary, B. P. Fairfax, M. Georges, B. T. Heijmans, M. Kähönen, Y. Kim, J. C. Knight, P. Kovacs, K. Krohn, S. Li, M. Loeffler, U. M. Marigorta, H. Mei, Y. Momozawa, M. Müller-Nurasyid, M. Nauck, M. Nivard, B. Penninx, J. Pritchard, O. Raitakari, O. Rotzchke, E. P. Slagboom, C. D. A. Stehouwer, M. Stumvoll, P. Sullivan, P. A. C. ‘t Hoen, J. Thiery, A. Tönjes, J. van Dongen, M. van Iterson, J. Veldink, U. Völker, C. Wijmenga, M. Swertz, A. Andiappan, G. W. Montgomery, S. Ripatti, M. Perola, Z. Kutalik, E. Dermitzakis, S. Bergmann, T. Frayling, J. van Meurs, H. Prokisch, H. Ahsan, B. Pierce, T. Lehtimäki, D. Boomsma, B. M. Psaty, S. A. Gharib, P. Awadalla, L. Milani, W. Ouwehand, K. Downes, O. Stegle, A. Battle, J. Yang, P. M. Visscher, M. Scholz, G. Gibson, T. Esko, L. Franke, Unraveling the polygenic architecture of complex traits using blood eQTL metaanalysis. Genomics (2018), p. 228.

14. T. Qi, Y. Wu, J. Zeng, F. Zhang, A. Xue, L. Jiang, Z. Zhu, K. Kemper, L. Yengo, Z. Zheng, eQTLGen Consortium, R. E. Marioni, G. W. Montgomery, I. J. Deary, N. R. Wray, P. M. Visscher, A. F. McRae, J. Yang, Identifying gene targets for brain-related traits using transcriptomic and methylomic data from blood. Nat. Commun. 9, 2282 (2018).

15. GTEx Consortium, Laboratory, Data Analysis &Coordinating Center (LDACC)—Analysis Working Group, Statistical Methods groups—Analysis Working Group, Enhancing GTEx (eGTEx) groups, NIH Common Fund, NIH/NCI, NIH/NHGRI, NIH/NIMH, NIH/NIDA, Biospecimen Collection Source Site—NDRI, Biospecimen Collection Source Site—RPCI, Biospecimen Core Resource— VARI, Brain Bank Repository—University of Miami Brain Endowment Bank, Leidos Biomedical— Project Management, ELSI Study, Genome Browser Data Integration &Visualization—EBI, Genome Browser Data Integration &Visualization—UCSC Genomics Institute, University of California Santa Cruz, Lead analysts:, Laboratory, Data Analysis &Coordinating Center (LDACC):, NIH program management:, Biospecimen collection:, Pathology:, eQTL manuscript working group:, A. Battle, C. D. Brown, B. E. Engelhardt, S. B. Montgomery, Genetic effects on gene expression across human tissues. Nature. 550, 204–213 (2017).

16. M. Fromer, P. Roussos, S. K. Sieberts, J. S. Johnson, D. H. Kavanagh, T. M. Perumal, D. M. Ruderfer, E. C. Oh, A. Topol, H. R. Shah, L. L. Klei, R. Kramer, D. Pinto, Z. H. Gümüş, A. E. Cicek, K. K. Dang, A. Browne, C. Lu, L. Xie, B. Readhead, E. A. Stahl, J. Xiao, M. Parvizi, T. Hamamsy, J. F. Fullard, Y.-C. Wang, M. C. Mahajan, J. M. J. Derry, J. T. Dudley, S. E. Hemby, B. A. Logsdon, K. Talbot, T. Raj, D. A. Bennett, P. L. De Jager, J. Zhu, B. Zhang, P. F. Sullivan, A. Chess, S. M. Purcell, L. A. Shinobu, L. M. Mangravite, H. Toyoshiba, R. E. Gur, C.-G. Hahn, D. A. Lewis, V. Haroutunian, M. A. Peters, B. K. Lipska, J. D. Buxbaum, E. E. Schadt, K. Hirai, K. Roeder, K. J. Brennand, N. Katsanis, E. Domenici, B. Devlin, P. Sklar, Gene expression elucidates functional impact of polygenic risk for schizophrenia. Nat. Neurosci. 19, 1442–1453 (2016).

17. B. Ng, C. C. White, H.-U. Klein, S. K. Sieberts, C. McCabe, E. Patrick, J. Xu, L. Yu, C. Gaiteri, D. A. Bennett, S. Mostafavi, P. L. De Jager, An xQTL map integrates the genetic architecture of the human brain’s transcriptome and epigenome. Nat. Neurosci. 20, 1418–1426 (2017).

18. N. Habib, I. Avraham-Davidi, A. Basu, T. Burks, K. Shekhar, M. Hofree, S. R. Choudhury, F. Aguet, E. Gelfand, K. Ardlie, D. A. Weitz, O. Rozenblatt-Rosen, F. Zhang, A. Regev, Massively parallel single-nucleus RNA-seq with DroNc-seq. Nat. Methods. 14, 955–958 (2017).

19. K. Burk, R. J. Pasterkamp, Disrupted neuronal trafficking in amyotrophic lateral sclerosis. Acta Neuropathol. 137, 859–877 (2019).

20. A. Brockington, P. R. Heath, H. Holden, P. Kasher, F. L. P. Bender, F. Claes, D. Lambrechts, M. Sendtner, P. Carmeliet, P. J. Shaw, Downregulation of genes with a function in axon outgrowth and synapse formation in motor neurones of the VEGFd/d mouse model of amyotrophic lateral sclerosis. BMC Genomics. 11 (2010), p. 203.

21. N. Hirokawa, R. Takemura, Molecular motors and mechanisms of directional transport in neurons. Nat. Rev. Neurosci. 6, 201–214 (2005).

22. D. Do-Ha, Y. Buskila, L. Ooi, Impairments in Motor Neurons, Interneurons and Astrocytes Contribute to Hyperexcitability in ALS: Underlying Mechanisms and Paths to Therapy. Mol. Neurobiol. 55, 1410–1418 (2018).

23. C. J. Donnelly, P.-W. Zhang, J. T. Pham, A. R. Haeusler, N. A. Mistry, S. Vidensky, E. L. Daley, E. M. Poth, B. Hoover, D. M. Fines, N. Maragakis, P. J. Tienari, L. Petrucelli, B. J. Traynor, J. Wang, F. Rigo, C. F. Bennett, S. Blackshaw, R. Sattler, J. D. Rothstein, RNA toxicity from the ALS/FTD C9ORF72 expansion is mitigated by antisense intervention. Neuron. 80, 415–428 (2013).

24. W. Xu, J. Xu, C9orf72 Dipeptide Repeats Cause Selective Neurodegeneration and Cell-Autonomous Excitotoxicity in Drosophila Glutamatergic Neurons. The Journal of Neuroscience. 38 (2018), pp. 7741–7752.

25. A. J. Waite, D. Bäumer, S. East, J. Neal, H. R. Morris, O. Ansorge, D. J. Blake, Reduced C9orf72 protein levels in frontal cortex of amyotrophic lateral sclerosis and frontotemporal degeneration brain with the C9ORF72 hexanucleotide repeat expansion. Neurobiology of Aging. 35 (2014), pp. 1779.e5–1779.e13.

26. W. Y. Ho, Y. K. Tai, J.-C. Chang, J. Liang, S.-H. Tyan, S. Chen, J.-L. Guan, H. Zhou, H.-M. Shen, E. Koo, S.-C. Ling, The ALS-FTD-linked gene product, C9orf72, regulates neuronal morphogenesis via autophagy. Autophagy. 15, 827–842 (2019).

27. L. Ferraiuolo, K. Meyer, T. W. Sherwood, J. Vick, S. Likhite, A. Frakes, C. J. Miranda, L. Braun, P. Heath, R. Pineda, C. E. Beattie, P. J. Shaw, C. C. Askwith, D. McTigue, B. K. Kaspar, Oligodendrocytes contribute to motor neuron death in ALS via SOD1-dependent mechanism. Proc. Natl. Acad. Sci. U. S. A. 113, E6496–E6505 (2016).

28. H. P. Nguyen, C. Van Broeckhoven, J. van der Zee, ALS Genes in the Genomic Era and their Implications for FTD. Trends Genet. 34, 404–423 (2018).

29. P. Khatri, M. Sirota, A. J. Butte, Ten years of pathway analysis: current approaches and outstanding challenges. PLoS Comput. Biol. 8, e1002375 (2012).

30. A. Subramanian, P. Tamayo, V. K. Mootha, S. Mukherjee, B. L. Ebert, M. A. Gillette, A. Paulovich, L. Pomeroy, T. R. Golub, E. S. Lander, J. P. Mesirov, Gene set enrichment analysis: A knowledge-based approach for interpreting genome-wide expression profiles. Proceedings of the National Academy of Sciences. 102, 15545–15550 (2005).

31. A. Liberzon, A. Subramanian, R. Pinchback, H. Thorvaldsdóttir, P. Tamayo, J. P. Mesirov, Molecular signatures database (MSigDB) 3.0. Bioinformatics. 27, 1739–1740 (2011).

32. J. Euesden, C. M. Lewis, P. F. O’Reilly, PRSice: Polygenic Risk Score software. Bioinformatics. 31, 1466–1468 (2014).

33. P. Mehta, W. Kaye, J. Raymond, R. Punjani, T. Larson, J. Cohen, O. Muravov, K. Horton, Prevalence of Amyotrophic Lateral Sclerosis — United States, 2015. MMWR. Morbidity and Mortality Weekly Report. 67 (2018), pp. 1285–1289.

34. S. Purcell, B. Neale, K. Todd-Brown, L. Thomas, M. A. R. Ferreira, D. Bender, J. Maller, P. Sklar, P. I. W. de Bakker, M. J. Daly, P. C. Sham, PLINK: a tool set for whole-genome association and population-based linkage analyses. Am. J. Hum. Genet. 81, 559–575 (2007).

35. G. Yu, F. Li, Y. Qin, X. Bo, Y. Wu, S. Wang, GOSemSim: an R package for measuring semantic similarity among GO terms and gene products. Bioinformatics. 26 (2010), pp. 976–978.

36. A. Battle, S. Mostafavi, X. Zhu, J. B. Potash, M. M. Weissman, C. McCormick, C. D. Haudenschild, K. B. Beckman, J. Shi, R. Mei, A. E. Urban, S. B. Montgomery, D. F. Levinson, D. Koller, Characterizing the genetic basis of transcriptome diversity through RNA-sequencing of 922 individuals. Genome Res. 24, 14–24 (2014).

37. Z. Zhu, F. Zhang, H. Hu, A. Bakshi, M. R. Robinson, J. E. Powell, G. W. Montgomery, M. E. Goddard, N. R. Wray, P. M. Visscher, J. Yang, Integration of summary data from GWAS and eQTL studies predicts complex trait gene targets. Nature Genetics. 48 (2016), pp. 481–487.

38. C. Hafemeister, R. Satija, Normalization and variance stabilization of single-cell RNA-seq data using regularized negative binomial regression. Genome Biol. 20, 296 (2019).

39. T. Stuart, A. Butler, P. Hoffman, C. Hafemeister, E. Papalexi, W. M. Mauck 3rd, Y. Hao, M. Stoeckius, P. Smibert, R. Satija, Comprehensive Integration of Single-Cell Data. Cel l. 177, 1888–1902.e21 (2019).

40. A. Butler, P. Hoffman, P. Smibert, E. Papalexi, R. Satija, Integrating single-cell transcriptomic data across different conditions, technologies, and species. Nat. Biotechnol. 36, 411–420 (2018).

41. R. D. Hodge, T. E. Bakken, J. A. Miller, K. A. Smith, E. R. Barkan, L. T. Graybuck, J. L. Close, B. Long, N. Johansen, O. Penn, Z. Yao, J. Eggermont, T. Höllt, B. P. Levi, S. I. Shehata, B. Aevermann, A. Beller, D. Bertagnolli, K. Brouner, T. Casper, C. Cobbs, R. Dalley, N. Dee, S.-L. Ding, R. G. Ellenbogen, O. Fong, E. Garren, J. Goldy, R. P. Gwinn, D. Hirschstein, C. D. Keene, M. Keshk, A. L. Ko, K. Lathia, A. Mahfouz, Z. Maltzer, M. McGraw, T. N. Nguyen, J. Nyhus, J. G. Ojemann, A. Oldre, S. Parry, S. Reynolds, C. Rimorin, N. V. Shapovalova, S. Somasundaram, A. Szafer, E. R. Thomsen, M. Tieu, G. Quon, R. H. Scheuermann, R. Yuste, S. M. Sunkin, B. Lelieveldt, D. Feng, L. Ng, A. Bernard, M. Hawrylycz, J. W. Phillips, B. Tasic, H. Zeng, A. R. Jones, C. Koch, E. S. Lein, Conserved cell types with divergent features in human versus mouse cortex. Nature. 573, 61–68 (2019).

42. B. B. Lake, R. Ai, G. E. Kaeser, N. S. Salathia, Y. C. Yung, R. Liu, A. Wildberg, D. Gao, H.-L. Fung, S. Chen, R. Vijayaraghavan, J. Wong, A. Chen, X. Sheng, F. Kaper, R. Shen, M. Ronaghi, J.-B. Fan, W. Wang, J. Chun, K. Zhang, Neuronal subtypes and diversity revealed by single-nucleus RNA sequencing of the human brain. Science. 352, 1586–1590 (2016).

43. GTEx Consortium, Human genomics. The Genotype-Tissue Expression (GTEx) pilot analysis: multitissue gene regulation in humans. Science. 348, 648–660 (2015).

44. F. Dudbridge, Power and predictive accuracy of polygenic risk scores. PLoS Genet. 9, e1003348 (2013).

45. S. Das, L. Forer, S. Schönherr, C. Sidore, A. E. Locke, A. Kwong, S. I. Vrieze, E. Y. Chew, S. Levy, M. McGue, D. Schlessinger, D. Stambolian, P.-R. Loh, W. G. Iacono, A. Swaroop, L. J. Scott, F. Cucca, F. Kronenberg, M. Boehnke, G. R. Abecasis, C. Fuchsberger, Next-generation genotype imputation service and methods. Nat. Genet. 48, 1284–1287 (2016).

46. S. McCarthy, S. Das, W. Kretzschmar, O. Delaneau, A. R. Wood, A. Teumer, H. M. Kang, C. Fuchsberger, P. Danecek, K. Sharp, Y. Luo, C. Sidore, A. Kwong, N. Timpson, S. Koskinen, S. Vrieze, L. J. Scott, H. Zhang, A. Mahajan, J. Veldink, U. Peters, C. Pato, C. M. van Duijn, C. E. Gillies, I. Gandin, M. Mezzavilla, A. Gilly, M. Cocca, M. Traglia, A. Angius, J. C. Barrett, D. Boomsma, K. Branham, G. Breen, C. M. Brummett, F. Busonero, H. Campbell, A. Chan, S. Chen, E. Chew, F. S. Collins, L. J. Corbin, G. D. Smith, G. Dedoussis, M. Dorr, A.-E. Farmaki, L. Ferrucci, L. Forer, R. M. Fraser, S. Gabriel, S. Levy, L. Groop, T. Harrison, A. Hattersley, O. L. Holmen, K. Hveem, M. Kretzler, J. C. Lee, M. McGue, T. Meitinger, D. Melzer, J. L. Min, K. L. Mohlke, J. B. Vincent, M. Nauck, D. Nickerson, A. Palotie, M. Pato, N. Pirastu, M. McInnis, J. B. Richards, C. Sala, V. Salomaa, D. Schlessinger, S. Schoenherr, P. E. Slagboom, K. Small, T. Spector, D. Stambolian, M. Tuke, J. Tuomilehto, L. H. Van den Berg, W. Van Rheenen, U. Volker, C. Wijmenga, D. Toniolo, E. Zeggini, P. Gasparini, M. G. Sampson, J. F. Wilson, T. Frayling, P. I. W. de Bakker, M. A. Swertz, S. McCarroll, C. Kooperberg, A. Dekker, D. Altshuler, C. Willer, W. Iacono, S. Ripatti, N. Soranzo, K. Walter, A. Swaroop, F. Cucca, C. A. Anderson, R. M. Myers, M. Boehnke, M. I. McCarthy, R. Durbin, Haplotype Reference Consortium, A reference panel of 64,976 haplotypes for genotype imputation. Nat. Genet. 48, 1279–1283 (2016).

47. A. E. Renton, E. Majounie, A. Waite, J. Simón-Sánchez, S. Rollinson, J. R. Gibbs, J. C. Schymick, H. Laaksovirta, J. C. van Swieten, L. Myllykangas, H. Kalimo, A. Paetau, Y. Abramzon, A. M. Remes, A. Kaganovich, S. W. Scholz, J. Duckworth, J. Ding, D. W. Harmer, D. G. Hernandez, J. O. Johnson, K. Mok, M. Ryten, D. Trabzuni, R. J. Guerreiro, R. W. Orrell, J. Neal, A. Murray, J. Pearson, I. E. Jansen, D. Sondervan, H. Seelaar, D. Blake, K. Young, N. Halliwell, J. B. Callister, G. Toulson, A. Richardson, A. Gerhard, J. Snowden, D. Mann, D. Neary, M. A. Nalls, T. Peuralinna, L. Jansson, V.-M. Isoviita, A.-L. Kaivorinne, M. Hölttä-Vuori, E. Ikonen, R. Sulkava, M. Benatar, J. Wuu, A. Chiò, G. Restagno, G. Borghero, M. Sabatelli, ITALSGEN Consortium, D. Heckerman, E. Rogaeva, L. Zinman, J. D. Rothstein, M. Sendtner, C. Drepper, E. E. Eichler, C. Alkan, Z. Abdullaev, S. D. Pack, A. Dutra, E. Pak, J. Hardy, A. Singleton, N. M. Williams, P. Heutink, S. Pickering-Brown, H. R. Morris, P. J. Tienari, B. J. Traynor, A hexanucleotide repeat expansion in C9ORF72 is the cause of chromosome 9p21-linked ALS-FTD. Neuron. 72, 257–268 (2011).

48. H. Heberle, G. V. Meirelles, F. R. da Silva, G. P. Telles, R. Minghim, InteractiVenn: a web-based tool for the analysis of sets through Venn diagrams. BMC Bioinformatics. 16, 169 (2015).

