## Supplemental Material for "Genetic analysis of amyotrophic lateral sclerosis identifies contributing pathways and cell types"

Supplemental Materials and Methods.

Fig. S1. Collection of gene sets and pathways within the Molecular Signature Database (MSigDB) used in this study.

Fig. S2. SNP set enrichment analysis used to dissect the biological function of significant cellular component and molecular function terms.

Fig. S3. The phenotypic variance explained by polygenic risk score across thirteen human cell types of human hippocampus and cortex obtained by DroNc-seq (*18*).

Figure S4. The phenotypic variance explained by polygenic risk score across twenty-four mouse brain cell types (*10*).

Table S1. Pathways that were significantly associated with ALS based on polygenic risk score analysis in the training dataset.

Table S2. Pathways that were significantly associated with ALS based on polygenic risk score analysis in the replication dataset after including known ALS GWAS loci as covariates.

Table S3. Significant functional associations for the significant pathway genes assessed by two‐sample Mendelian randomization.

Table S4. Demographic description of the cohorts used in the training and replication datasets.

Consortia.

**Supplemental Materials and Methods**

Genome-wide genotyping

For the training/replication cohorts, case samples were genotyped in the Laboratory of Neurogenetics, National Institutes of Health using HumanOmniExpress (version 1.0 genotyping 716,503 SNPs) according to the manufacturer's protocol (Illumina Inc., San Diego, CA). The remainder US control cases had been previously genotyped on HumanOmni BeadChips (Illumina) as part of other GWAS efforts, and these data were downloaded from the dbGaP web (*7*). Additional data from the *HYPERGENES* project and the Wellcome Trust Case Control Consortium were included as Italian and UK controls (for demographic description of the cohorts used in this study, see [***SI Appendix,***](https://www.pnas.org/lookup/suppl/doi:10.1073/pnas.1918314117/-/DCSupplemental) **Table S4)**. Familial cases were included in the analysis.

Genotyping data quality control procedures and imputation

We applied standard quality-control procedures to our genotype data. Briefly, individuals with low call rates (< 95%), heterozygosity outliers (F-statistic cutoff of > −0.15 and < 0.15), and ancestry outliers (+ /− 6 standard deviations from means of eigenvectors 1 and 2 of the 1000 Genomes phase 3 CEU and TSI populations from principal components) were excluded. Further, for genotype QC, variants with a missingness rate of > 5%, exhibiting deviation from Hardy–Weinberg Equilibrium (HWE) in controls (p < 10^-6^) and palindromic SNPs were excluded. Cryptic relatedness was assessed using a Pi hat of more than 0.125. Accordingly, individuals who shared more than 12.5% of their genome were excluded from the analysis. The remaining samples were imputed using the Haplotype Reference Consortium (HRC) on the Michigan Imputation Server pipeline using Minimac4 (*45*) under default settings with Eagle v2.4 phasing based on *Haplotype Reference Consortium r1.1 2016* (*46*)([http://www.haplotype-reference-consortium.org](http://www.haplotype-reference-consortium.org/)). Samples from the United States, Italy, the United Kingdom, Belgium, and France were imputed as a single group, and variants with an imputation quality (R^2^ >0.8) were included. Those variants were additionally filtered post-imputation to exclude variants with minor allele frequency < 0.01 and a missingness rate of > 15%.

Repeat-Primed PCR for *C9orf72* expansions detection

Repeat-primed PCR was performed according to an established protocol (*47*). Briefly, 100 ng of genomic DNA was added to the PCR mixture. Fragment length analysis was performed on an ABI 3730xl genetic analyzer (Applied Biosystems, Foster City, CA, USA), and data were analyzed using GeneScan software (version 4, Applied Biosystems).

Venn diagram

Venn diagrams were generated using *InteractiVenn*, a web-based tool for the analysis of sets (*48*).

**Supplemental Figures**

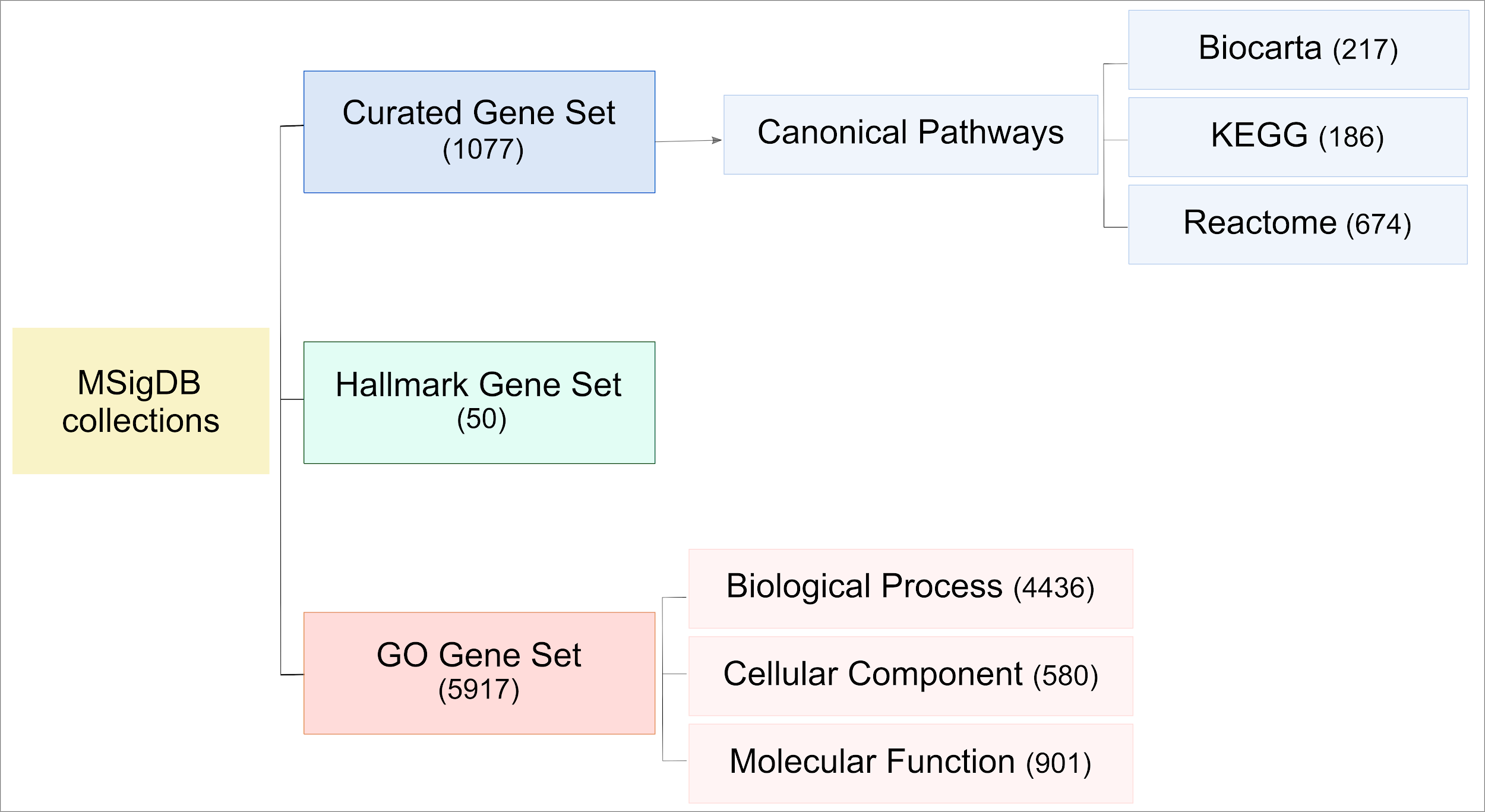

**Fig. S1. Collection of gene sets and pathways within the Molecular Signature Database (MSigDB) used in this study.**

The numbers in brackets represent the number of gene sets within each collection.

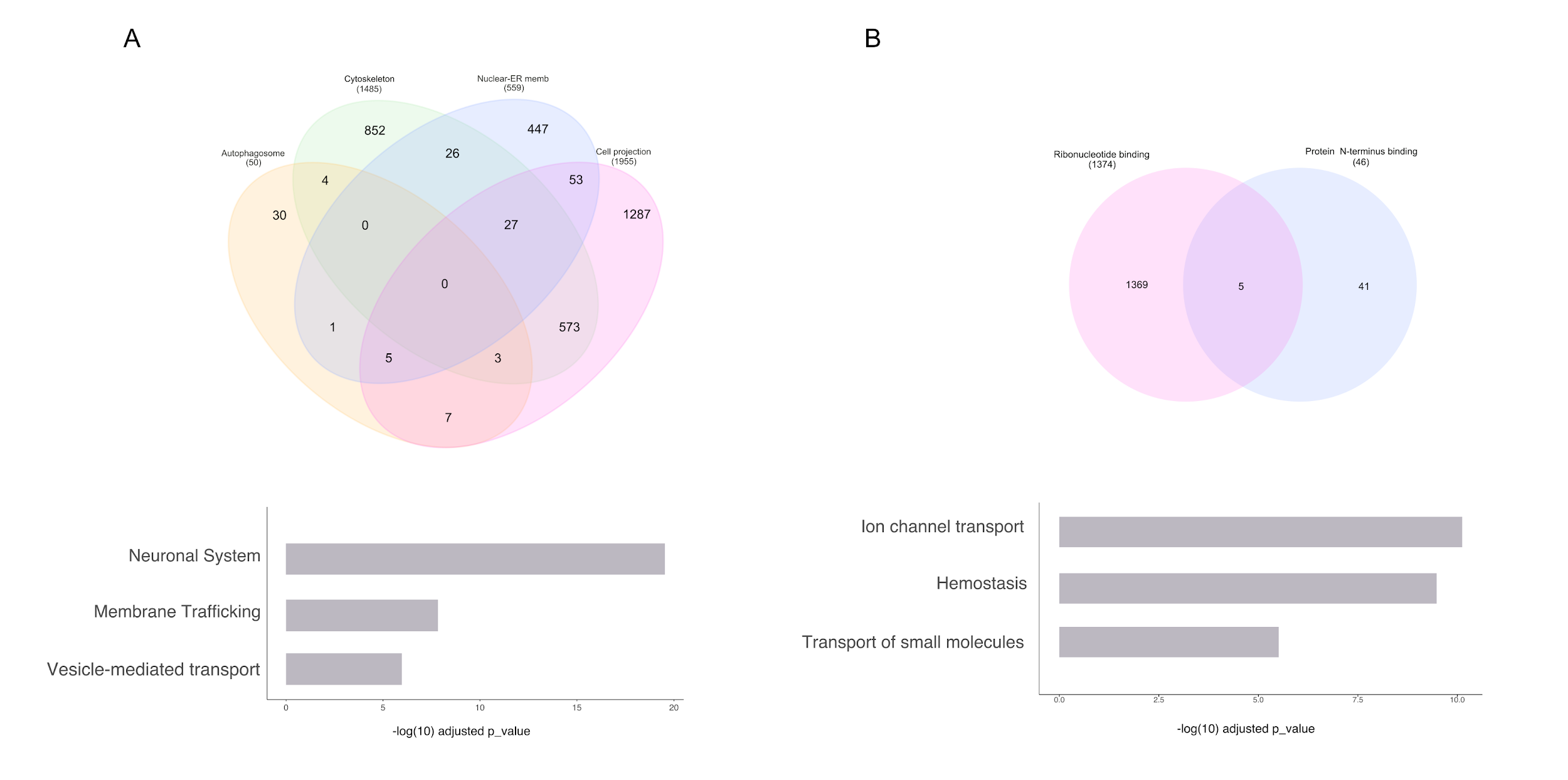

**Fig. S2. SNP set enrichment analysis used to dissect the biological function of significant cellular component and molecular function terms.**

The Venn diagrams in the upper panel show the number of SNPs in each term and the overlap between terms. The lower panels show the top three enriched pathways in (A) cellular component and (B) molecular function.

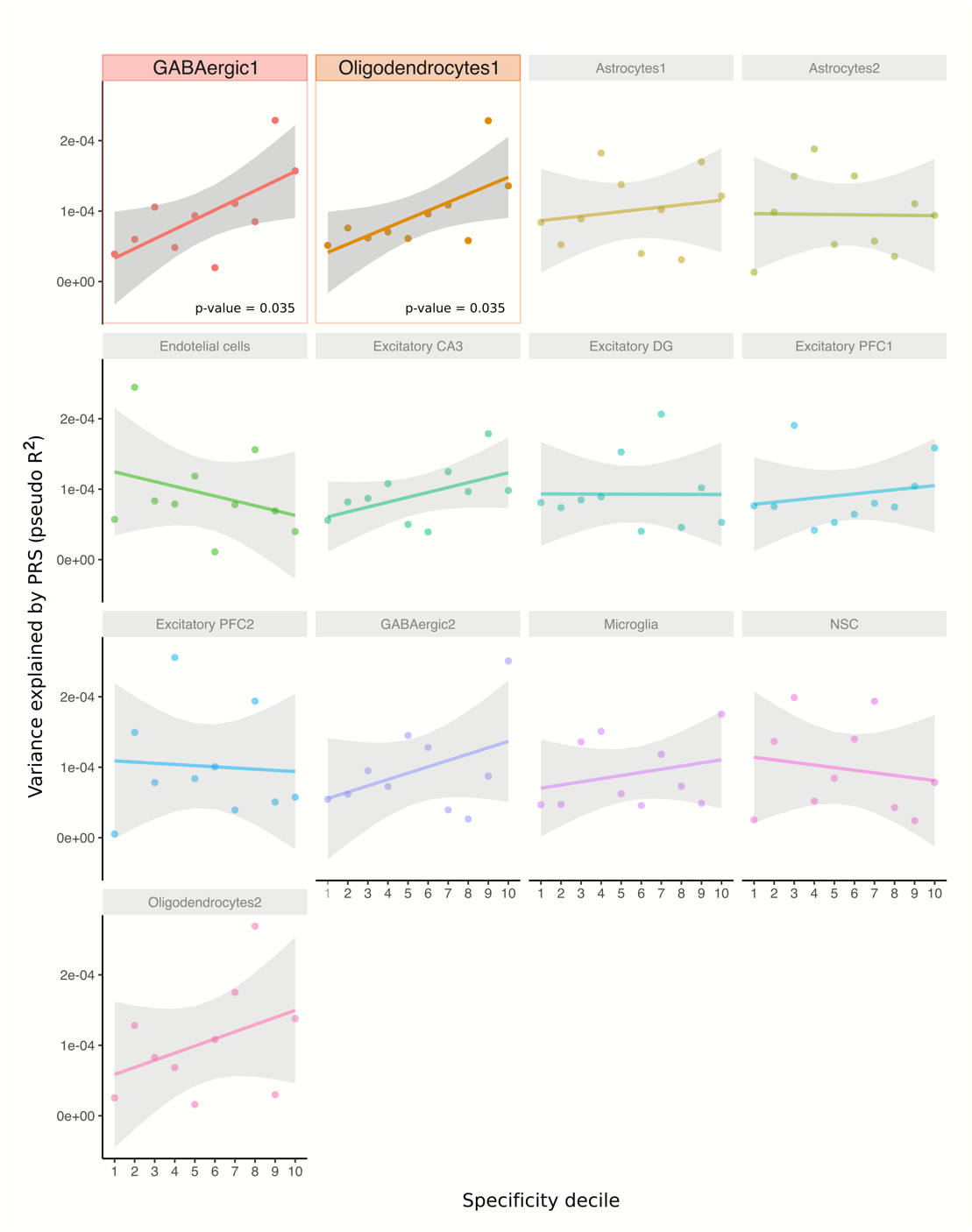

**Fig. S3. The phenotypic variance explained by polygenic risk score across thirteen human cell types of human hippocampus and cortex obtained by DroNc-seq** **(*18*)****.**

The y-axis corresponds to the phenotypic variance explained by the polygenic risk score (pseudo-R^2^), and the x-axis depicts deciles 1 to 10. The color pictures show the significant cell types and the significant p-values of the linear regression fit models. In contrast, the semitransparent images show the cell types not significantly associated with the disease. The regression line depicts the association between PRS.R2 (pseudo R2, adjusted by prevalence) and the specificity decile in each cell type. The grey shading shows the 95% confidence interval of the regression model. DG., dentate gyrus; NCS, neural stem cell; PFC., prefrontal cortex.

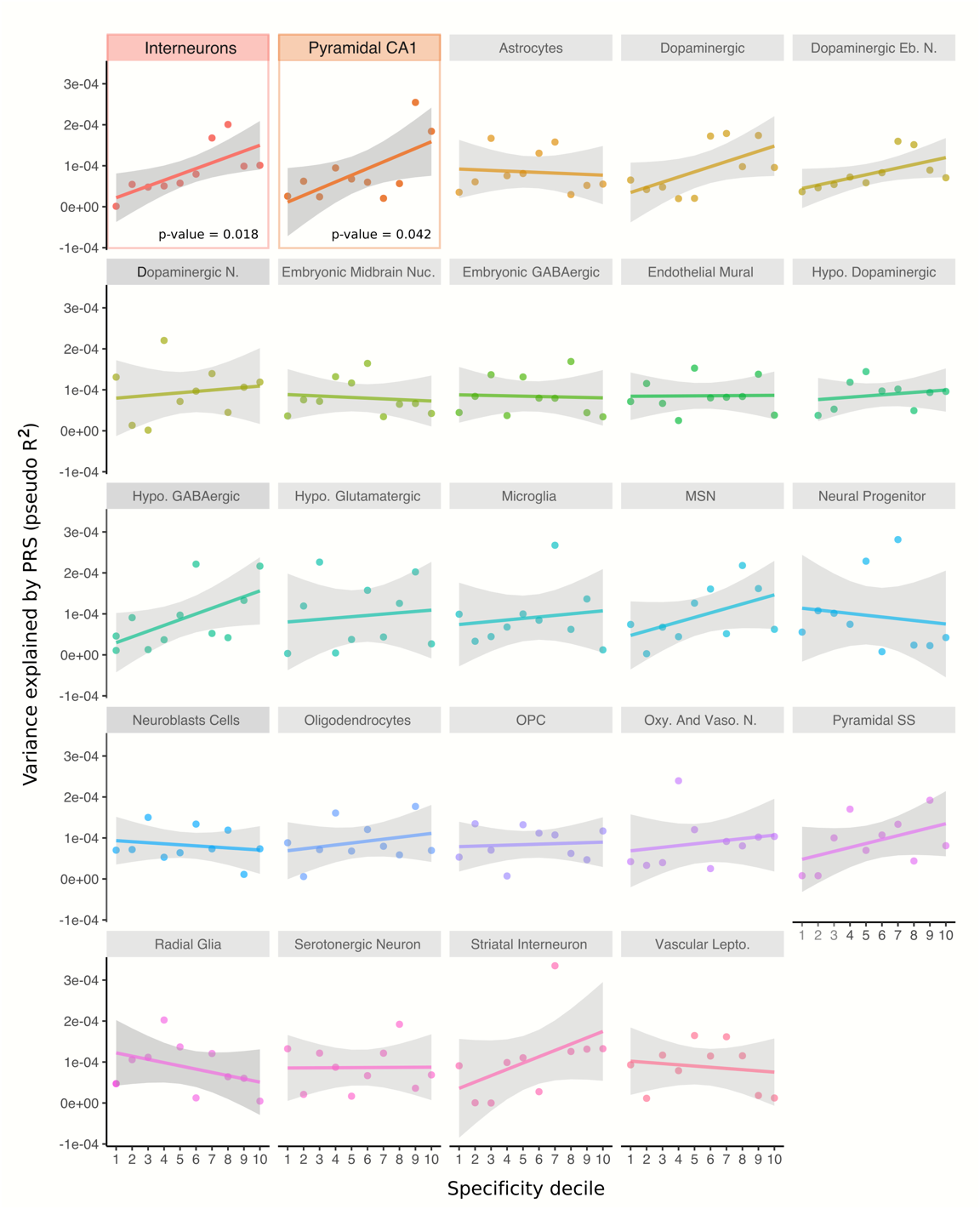

**Figure S4. The phenotypic variance explained by polygenic risk score across twenty-four mouse brain cell types (*10*).**

The y-axis corresponds to the phenotypic variance explained by the polygenic risk score (pseudo-R^2^), and the x-axis depicts deciles 1 to 10. The color pictures show the significant cell types and the significant p-values of the linear regression fit models. In contrast, the semitransparent images show the cell types not significantly associated with the disease. The regression line depicts the association between PRS.R2 (pseudo R2, adjusted by prevalence) and the specificity decile in each cell type. Cort., cortical; Eb. N., embryonic nucleus; Emb., embryonic; Hypo., hypothalamic; Lepto., leptomeningeal; MSN, medium spiny neurons; N., neuron; Oxy., oxytocin; P., progenitor; and Vaso., vasopressin.

**Supplementary Tables**

**Table S1. Pathways that were significantly associated with ALS based on polygenic risk score analysis in the training dataset.** SE, Standard Error; FDR, False Discovery Rate.

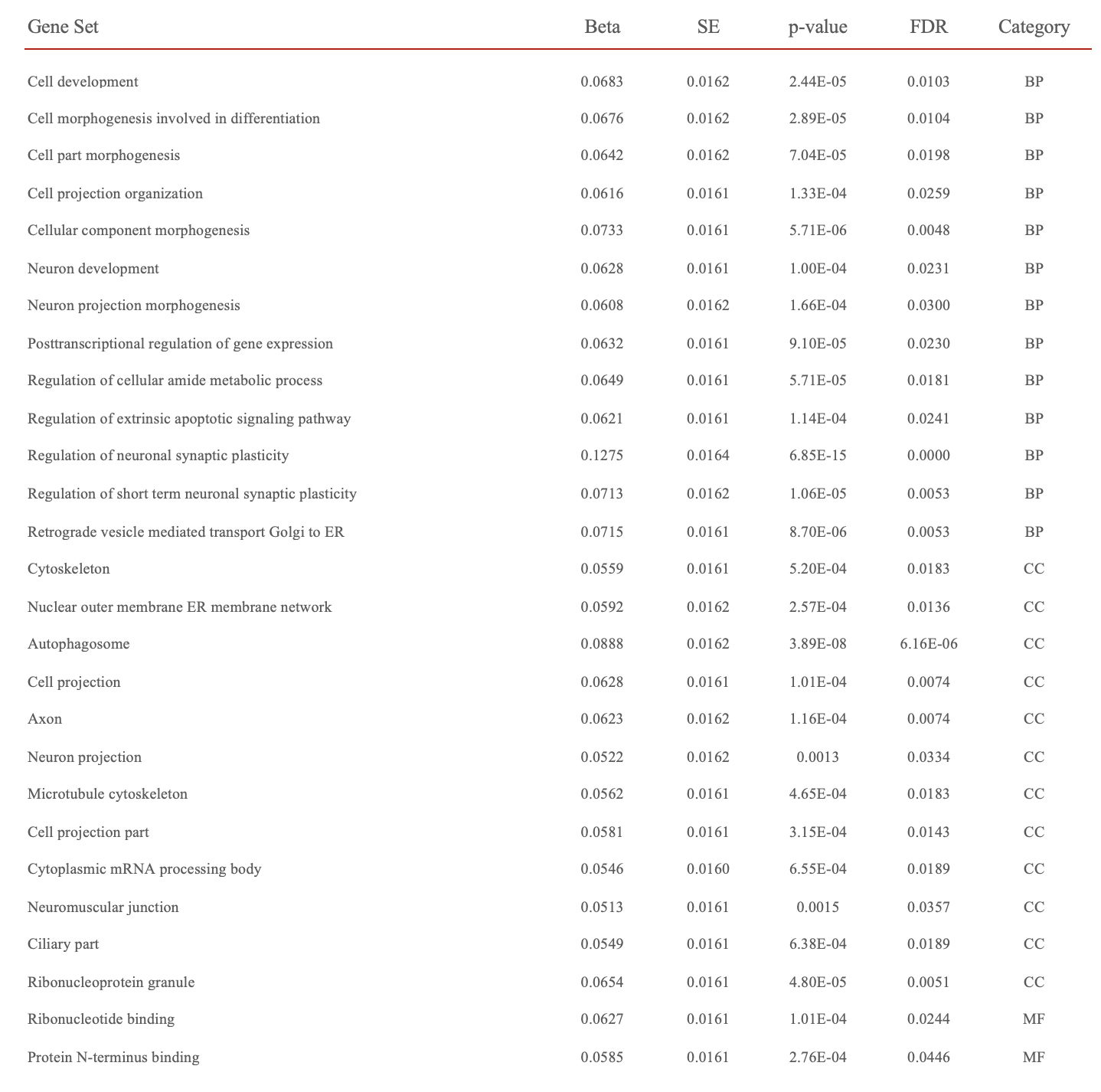

**Table S2. Pathways that were significantly associated with ALS based on polygenic risk score analysis in the replication dataset after including known ALS GWAS loci as covariates.** SE, Standard Error.

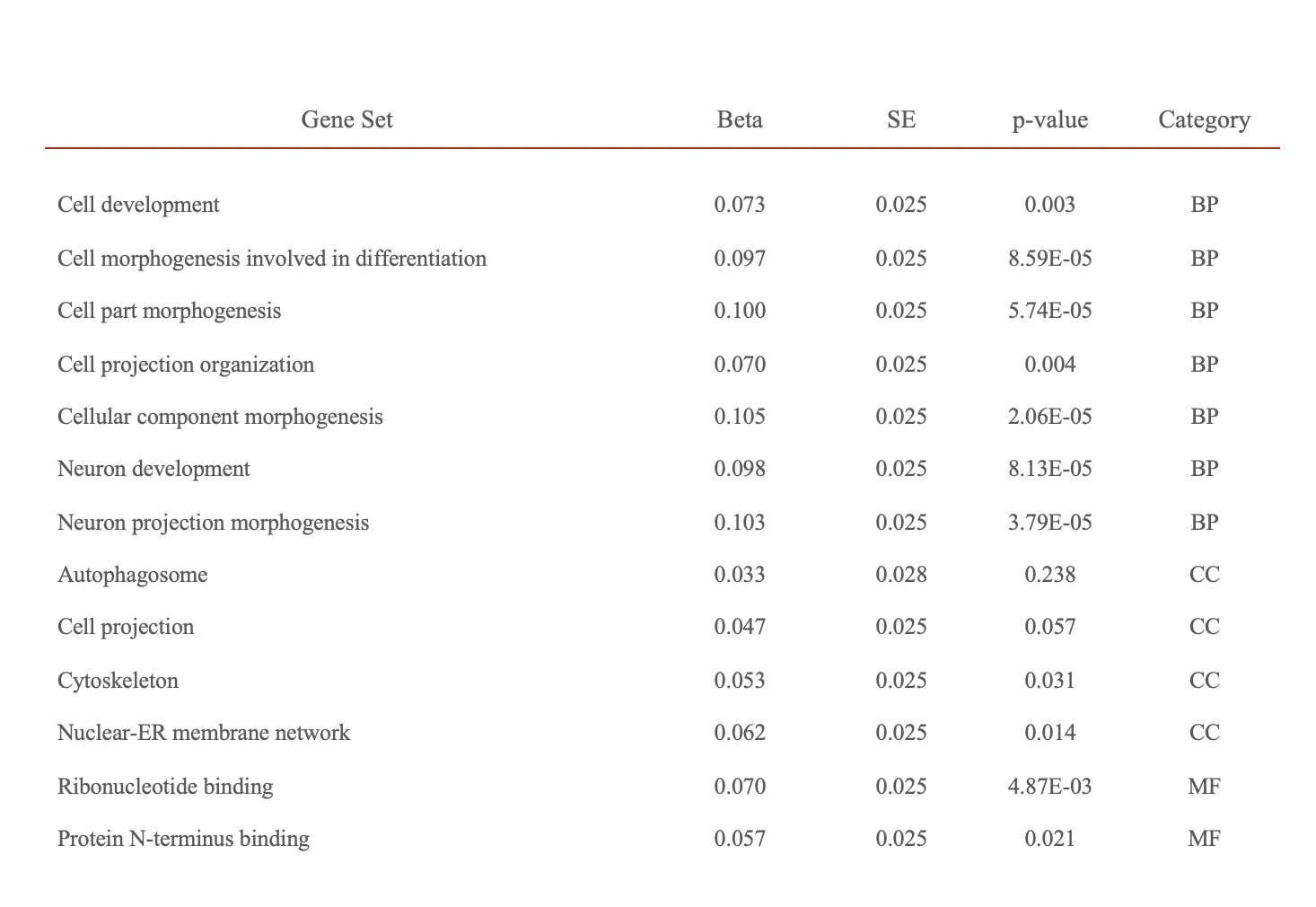

**Table S3. Significant functional associations for the significant pathway genes assessed by two‐sample Mendelian randomization.**

FDR, adjusted false discovery rate at single SNP p-value after performing Mendelian randomization between eQTL resources (exposures) versus the Nicolas GWAS (outcome). SE, standard error. p_HEIDI, p-value HEIDI.

| Gene | SNP | Beta | SE | p-value | FDR | p_HEIDI | Analyte |
| --- | --- | --- | --- | --- | --- | --- | --- |
| SCFD1 | rs7144204 | -0.309 | 0.063 | 8.48E-07 | 0.001 | 0.381 | Blood |
| PLXNB2 | rs62241220 | -0.268 | 0.061 | 1.03E-05 | 0.002 | 0.362 | Blood |
| MAPKAPK3 | rs13096264 | -0.123 | 0.034 | 2.67E-04 | 0.019 | 0.018 | Blood |
| ACSL5 | rs10885342 | 0.087 | 0.023 | 1.43E-04 | 0.047 | 0.197 | Blood |
| ATG16L2 | rs2282613 | 0.569 | 0.184 | 0.002 | 0.039 | 0.050 | Blood |
| SCFD1 | rs2070339 | 0.200 | 0.045 | 7.55E-06 | 0.003 | 0.340 | Brain |

**Table S4. Demographic description of the cohorts used in the training and replication datasets.**

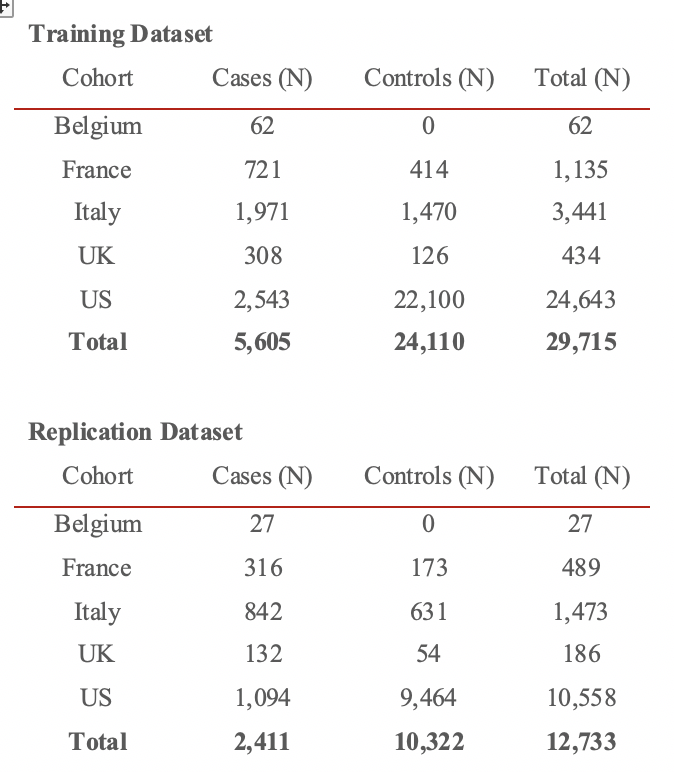

**Consortia**

**The members of the ITALSGEN Consortium are:**

Francesco Pio Ausiello^1^, Marco Barberis^2^, Ilaria Bartolomei^3^, Stefania Battistini^4^, Michele Benigni^4^, Enrica Bersano^5^, Giuseppe Borghero^6^, Maura Brunetti^7^, Andrea Calvo^2^, Antonino Cannas^6^, Antonio Canosa^2^, Margherita Capasso^8^, Claudia Caponnetto^9^, Patrizio Cardinali^10^, Paola Carrera^11^, Federico Casale^2^, Adriano Chiò^2^, Tiziana Colletti^12^, Francesca L Conforti^13^, Amelia Conte^14^, Elisa Conti^15^, Massimo Corbo^16^, Eleonora Dalla Bella^5^, Giovanni Defazio^6^, Raffaele Dubbioso^1^, Antonio Fasano^17^, Cinzia Femiano^18^, Carlo Ferrarese^15^, Nicola Fini^17^, Gianluca Floris^6^, Giuseppe Fuda^2^, Fabio Giannini^4^, Carlo Guidi^19^, Antonio Ilardi^2^, Vincenzo La Bella^12^, Serena Lattante^14^, Giuseppe Lauria^5,20^, Giancarlo Logroscino^21^, Francesco O Logullo^22^, Christian Lunetta^23^, Gianluigi Mancardi^9^, Paola Mandich^9^, Jessica Mandrioli^17^, Umberto Manera^2^, Giuseppe Marangi^14^, Kalliopi Marinou^24^, Maria Giovanna Marrosu^6^, Maurizio Melis^6^, Sonia Messina^25^, Cristina Moglia^2^, Maria Rosaria Monsurrò^18^, Gabriele Mora^26^, Lorena Mosca^27^, Maria Rita Murru^6^, Patrizia Occhineri^28^, Paola Origone^9^, Antonio Petrucci^29^, Giovanni Piccirillo^18^, Angelo Pirisi^28^, Maura Pugliatti^28^, Gabriella Restagno^7^, Claudia Ricci^4^, Nilo Riva^11^, Massimo Russo^30^, Mario Sabatelli^14^, Gessica Sala^15^, Fabrizio Salvi^3^, Marialuisa Santarelli^31^, Lucio Santoro^1^, Riccardo Sideri^24^, Isabella Simone^21^, Rossella Spataro^12^, Raffaella Tanel^32^, Gioacchino Tedeschi^18^, Anna Ticca^33^, Antonella Torriello^34^, Maria Claudia Torrieri^2^, Lucio Tremolizzo^15^, Francesca Trojsi^18^, Rosario Vasta^2^, Giuseppe Vita^25^, Paolo Volanti^35^, Marcella Zollino^14,20^

1. Università degli Studi di Napoli Federico II, Naples, Italy
2. Rita Levi Montalcini' Department of Neuroscience, Amyotrophic Lateral Sclerosis Center, University of Turin, Turin, Italy
3. Center for Diagnosis and Cure of Rare Diseases, Department of Neurology, IRCCS Institute of Neurological Sciences, Bologna, Italy
4. Department of Medical, Surgical and Neurological Sciences, University of Siena, Siena, Italy
5. Neurology and Headache Unit, Fondazione IRCCS Istituto Neurologico "Carlo Besta”, Milan, Italy
6. Department of Neurology, Azienda Universitario Ospedaliera di Cagliari and University of Cagliari, Cagliari, Italy
7. Molecular Genetics Unit, Department of Clinical Pathology, A.S.O. O.I.R.M.-S. Anna, 10126 Turin, Italy
8. Department of Neurology, University of Chieti, Chieti, Italy
9. Department of Neurosciences, Ophthalmology, Genetics, Rehabilitation, Maternal and Child Health, IRCCS Azienda Ospedaliero-Universitaria San Martino IST, Genoa, Italy
10. Amministrazione ASUR Zona Territoriale 11, Fermo, Italy
11. Department of Neurology and Institute of Experimental Neurology (INSPE), IRCCS San Raffaele Scientific Institute, Milan, Italy
12. ALS Clinical Research Center, Bio. Ne. C., University of Palermo, Palermo, Italy
13. Institute of Neurological Sciences, National Research Council, Mangone, Cosenza, Italy
14. Centro Clinico NEMO-Roma, Neurological Institute, Catholic University and I.C.O.M.M. Association for ALS Research, Rome, Italy
15. Neurology Unit, School of Medicine and Surgery and NeuroMI, University of Milano-Bicocca, Monza, Italy
16. Department of Neurorehabilitation Sciences (P.T., M.C.), Casa Cura Policlinico, Milan, Italy
17. Department of Neuroscience, S. Agostino-Estense Hospital, University of Modena and Reggio Emilia, Modena, Italy
18. Department of Medical, Surgical Neurological Metabolic and Aging Sciences, Second University of Naples, Naples, Italy
19. AUSL della Romagna, Forlì, Italy
20. Department of Biomedical and Clinical Sciences "Luigi Sacco", University of Milan, Milan, Italy
21. Department of Basic Medical Sciences, Neurosciences and Sense Organs, University of Bari, Bari, Italy
22. Neurological Clinic, Marche Polytechnic University, Ancona, Italy
23. NeuroMuscular Omnicenter, Serena Onlus Foundation, Milan, Italy
24. Department of Neurological Rehabilitation, Fondazione Salvatore Maugeri, IRCCS, Istituto Scientifico di Milano, Milan, Italy
25. OUC Neurology and Neuromuscular Disorders, University of Messina, Italy
26. ALS Center, ICS Maugeri, IRCCS, Milan, Italy
27. Department of Laboratory Medicine, Medical Genetics, Niguarda Ca' Granda Hospital, Milan, Italy
28. Department of Biomedical and Surgical Sciences, Section of Neurological, Psychiatric and Psychological Sciences, University of Ferrara, Ferrara, Italy
29. Neurology Department, San Camillo Hospital, Rome, Italy
30. Centro Clinico NEMO-Messina, Messina, Italy
31. Department of Medicine, Azienda Complesso Ospedaliero, San Filippo Neri, Rome, Italy
32. Department of Neurology, Santa Chiara Hospital, Trento, Italy
33. Department of Neurology, Azienda Ospedaliera San Francesco, Nuoro, Italy
34. AOU OO.RR. San Giovanni di Dio Ruggi d'Aragona Salerno, Salerno, Italy
35. Neurorehabilitation Unit/ALS Center, Scientific Clinical Institutes (ICS) Maugeri, IRCCS, Mistretta, Messina, Italy

**The members of the International ALS Genomics Consortium are:**

Yevgeniya Abramzon^1,2^, Sampath Arepalli^3^, Robert H. Baloh^4^, Robert Bowser^5^, Christopher B. Brady^6^, Alexis Brice^7,8^, James Broach^9^, Roy H. Campbell^10^, William Camu^11^, Ruth Chia^1^, Adriano Chiò^12,13^, John Cooper-Knock^14^, Daniele Cusi^15^, Jinhui Ding^16^, Carsten Drepper^18^, Vivian E. Drory^19^, Travis L. Dunckley^20^, John D. Eicher^21^, Faraz Faghri^22,10^, Eva Feldman^23^, Mary Kay Floeter^24^, Pietro Fratta^2^, Joshua T. Geiger^25^, Glenn Gerhard^20^, J. Raphael Gibbs^16^, Summer B. Gibson^26^, Jonathan D. Glass^27^, Stephen A. Goutman^23^, John Hardy^28^, Matthew B. Harms^29^, Terry D. Heiman-Patterson^30,31^, Dena G. Hernandez^3^, Lilja Jansson^32^, Freya Kamel^33^, Janine Kirby^15^, Neil W. Kowall^34^, Hannu Laaksovirta^32^, John E. Landers^35^, Francesco Landi^36^, Isabelle Le Ber^7,8^, Serge Lumbroso^37^, Daniel JL. MacGowan^38^, Nicholas J. Maragakis^39^, Gabriele Mora^40^, Kevin Mouzat^37^, Natalie A. Murphy^1^, Liisa Myllykangas^41^, Mike A. Nalls^22,42^, Richard W. Orrell^43^, Lyle W. Ostrow^39^, Roger Pamphlett^44^, Stuart Pickering-Brown^45^, Erik Pioro^46^, Hannah A. Pliner^1^, Stefan M. Pulst^26^, John M. Ravits^47^, Alan E. Renton^1,48^, Alberto Rivera^1^, Wim Robberecht^49^, Ekaterina Rogaeva^50^, Sara Rollinson^45^, Jeffrey D. Rothstein^39^, Erika Salvi^17^, Sonja W. Scholz^25,39^, Michael Sendtner^51^, Pamela J. Shaw^15^, Katie C. Sidle^28^, Zachary Simmons^52^, Andrew B. Singleton^22^, David J. Stone^21^, Pentti J. Tienari^32^, Bryan J. Traynor^1,39^, John Q. Trojanowski^53^, Juan C. Troncoso^54^, Miko Valori^32^, Philip Van Damme^49,55^, Vivianna M. Van Deerlin^53^, Ludo Van Den Bosch^49^, Lorne Zinman^56^

1. Neuromuscular Diseases Research Section, Laboratory of Neurogenetics, National Institute on Aging, Bethesda, MD 20892, USA
2. Sobell Department of Motor Neuroscience and Movement Disorders, Institute of Neurology, University College London, London, WC1N 3BG, UK
3. Genomics Technology Group, Laboratory of Neurogenetics, National Institute on Aging, Bethesda, MD 20892, USA
4. Department of Neurology, Cedars-Sinai Medical Center, Los Angeles, CA 90048, USA
5. Division of Neurology, Barrow Neurological Institute, Phoenix, AZ 85013, USA
6. Research and Development Service, Veterans Affairs Boston Healthcare System, Boston, MA 02130, USA
7. Centre de Recherche de l’Institut du Cerveau et de la Moelle épinière, Université Pierre et Marie Curie, Paris, France
8. INSERM U975, Paris, France
9. Department of Biochemistry, Penn State College of Medicine, Hershey, PA 17033, USA
10. Department of Computer Science, The University of Illinois at Urbana-Champaign, 201 North Goodwin Avenue, Urbana, IL 61801, USA
11. ALS reference center, Gui de Chauliac hospital, CHU and Univ Montpellier, Montpellier France
12. ‘Rita Levi Montalcini’ Department of Neuroscience, University of Turin, Via Verdi 8, Turin, 10124, Italy
13. Neuroscience Institute of Torino, University of Turin, Turin, 10124, Italy
14. Department of Neuroscience, University of Sheffield, Sheffield, S10 2HQ, UK
15. Bio4Dreams Scientific Unit Bio4Dreams - Business Nursery for Life Sciences Milano Italy16.
16. Computational Biology Core, Laboratory of Neurogenetics, National Institute on Aging, Bethesda, MD 20892, USA.
17. Neurology and Headache Unit, Fondazione IRCCS Istituto Neurologico "Carlo Besta”, Milan, Italy
18. Institute for Clinical Neurobiology, University of Würzburg, Würzburg, D-97078, Germany
19. Department of Neurology, Tel-Aviv Sourasky Medical Center, Tel-Aviv, Israel
20. Department of Pathology, Penn State College of Medicine, Hershey, PA 17033, USA
21. Genetics, Genetics and Pharmacogenomics, Merck Research Laboratories, Merck & Co., Inc., West Point, PA 19486, USA
22. Molecular Genetics Section, Laboratory of Neurogenetics, National Institute on Aging, Bethesda, MD 20892, USA
23. Department of Neurology, University of Michigan, 1500 E Medical Center Dr, Ann Arbor, MI 48109, USA
24. Motor Neuron Disorders Unit, Laboratory of Neurogenetics, National Institute of Neurological Disorders and Stroke, Bethesda, MD 20892, USA
25. Neurodegenerative Diseases Research Unit, Laboratory of Neurogenetics, National Institute of Neurological Disorders and Stroke, Bethesda, MD 20892, USA
26. Department of Neurology, University of Utah School of Medicine, 175 North Medical Drive East, Salt Lake City, UT 84132, USA
27. Department of Neurology, Emory University School of Medicine, Atlanta, GA 30322, USA
28. Department of Molecular Neuroscience and Reta Lila Weston Laboratories, Institute of Neurology, University College London, London, WC1N 3BG, UK
29. Department of Neurology, Columbia University, New York, NY 10032, USA
30. Department of Neurology, Drexel University College of Medicine, Philadelphia, PA 19102, USA
31. Department of Neurology, Temple University, 7602 Central Ave, Philadelphia, PA 19111, USA
32. Department of Neurology, University of Helsinki, Helsinki, FIN-02900, Finland
33. Epidemiology Branch, National Institute of Environmental Health Sciences, Durham, NC 27709, USA
34. Department of Neurology, Veterans Affairs Boston Healthcare System, Boston, MA 02130, USA
35. Department of Neurology, University of Massachusetts Medical School, Worcester, MA 01605, USA
36. Department of Geriatrics, Neurosciences and Orthopedics, Center for Geriatric Medicine, Catholic University of Sacred Heart, Rome, 00168, Italy
37. Service de Biochimie, CHU de Nîmes, Nîmes, France
38. New York Hospital Cornell University Medical Center 1305 York Avenue NYC NY 10021
39. Department of Neurology, Johns Hopkins University, Baltimore, MD 21287, USA
40. ALS Center, ICS Maugeri, IRCCS, Via Camaldoli, 64, Milan, 20138, Italy
41. Department of Pathology, Haartman Institute/HUSLAB, University of Helsinki and Folkhalsan Research Center (LM), Helsinki, FIN-02900, Finland
42. Data Tecnica International, Glen Echo, MD 20812, USA
43. Department of Clinical Neuroscience, Institute of Neurology, University College London, London, NW2 2PG, UK
44. Discipline of Pathology, Brain and Mind Centre, University of Sydney, Camperdown, NSW 2050, Australia
45. Faculty of Human and Medical Sciences, University of Manchester, Manchester, M13 9PT, UK
46. Department of Neurology, Cleveland Clinic, Cleveland, OH 44195, USA
47. Department of Neuroscience, Experimental Neurology and Leuven Research Institute for Neuroscience and Disease, University of California San Diego, 9500 Gilman Drive, La Jolla, CA 92093, USA
48. Department of Neuroscience, Ronald M. Loeb Center for Alzheimer's Disease, Icahn School of Medicine at Mount Sinai, New York, NY 10029, USA
49. Department of Neurosciences, Experimental Neurology and Leuven Research Institute for Neuroscience and Disease, University of Leuven, Leuven, 3000, Belgium
50. Division of Neurology, Tanz Centre for Research of Neurodegenerative Diseases and Toronto Western Hospital, University of Toronto, Toronto, M5S 3H2, Canada
51. Department of Neurology, Institute for Clinical Neurobiology, University of Würzburg, Würzburg, D-97078, Germany
52. Department of Neurology, Penn State College of Medicine, Hershey, PA 17033, USA
53. Department of Pathology and Laboratory Medicine, University of Pennsylvania, Philadelphia, PA 19104, USA
54. Clinical and Neuropathology Core, Johns Hopkins University, Baltimore, MD 21287, USA
55. VIB, Center for Brain & Disease Research, Laboratory of Neurobiology, University of Leuven, Leuven, 3000, Belgium
56. Division of Neurology, Sunnybrook Health Sciences Centre, University of Toronto, Toronto, M4N 3M5, Canada
